# A Highly Selective Response to Food in Human Visual Cortex Revealed by Hypothesis-Free Voxel Decomposition

**DOI:** 10.1101/2022.06.21.496922

**Authors:** Meenakshi Khosla, N Apurva Ratan Murty, Nancy Kanwisher

## Abstract

Prior work has identified cortical regions selectively responsive to specific categories of visual stimuli. However, this hypothesis-driven work cannot reveal how prominent these category selectivities are in the overall functional organization of visual cortex, or what others might exist that scientists have not thought to look for. Further, standard voxel-wise tests cannot detect distinct neural selectivities that coexist within voxels. To overcome these limitations, we used data-driven voxel decomposition methods to identify the main components underlying fMRI responses to thousands of complex photographic images (Allen et al 2021). Our hypothesis-neutral analysis rediscovered components selective for faces, places, bodies, and words, validating our method and showing that these selectivities are dominant features of the ventral visual pathway. The analysis also revealed an unexpected component with a distinct anatomical distribution that responded highly selectively to images of food. Alternative accounts based on low to mid-level visual features like color, shape or texture failed to account for the food selectivity of this component. High-throughput testing and control experiments with matched stimuli on a highly accurate computational model of this component confirm its selectivity for food. We registered our methods and hypotheses before replicating them on held-out participants and in a novel dataset. These findings demonstrate the power of data-driven methods, and show that the dominant neural responses of the ventral visual pathway include not only selectivities for faces, scenes, bodies, and words, but also the visually heterogeneous category of food, thus constraining accounts of when and why functional specialization arises in the cortex.

## Introduction

The last few decades of research in human cognitive neuroscience have revealed the functional organization of the cortex in rich detail. This organization features a set of regions that are selectively engaged in single mental processes, from perceiving faces or scenes or music, to understanding the meaning of a sentence, to inferring the content of another person’s thoughts. Why do our brains have these particular specializations, and apparently not others? To answer this question, we need a more complete inventory of human cortical specializations, one that reflects not just the idiosyncratic hypotheses scientists have already thought to test, but the actual functional organization of the cortex itself. Here, we tackle this question for the ventral visual pathway, by searching in a hypothesis-neutral fashion for the dominant neural response profiles in this region in a large recently-released public dataset of fMRI responses to thousands of natural images in each of 8 participants^1^.

Extensive evidence^2–7^ from neurological patients, fMRI, and intracranial recording and stimulation has demonstrated that the ventral visual pathway contains distinct regions causally engaged in the perception of faces, scenes, bodies, and words. But are these categories the main ones, or might others exist that have not yet been found? The current evidence does not answer this question for several reasons. First, prior research on the ventral pathway has tested a relatively small number of stimulus categories, which may not have subtended the relevant part of stimulus space preferred by some neural populations. Second, this work has proceeded in a largely hypothesis-driven fashion, and so may have missed neural populations with response profiles scientists have not thought to test. Third, prior research based on voxelwise contrasts is not well suited for discovering neural populations whose high selectivity is masked because the fMRI signal averages their responses with the responses of other neural populations cohabiting the same voxels^8^.

Here, we overcome these three limitations by analyzing fMRI responses to the very broad and large set of natural stimuli in the Natural Scenes Dataset (NSD^1^) with a data-driven analysis method that can de-mix the underlying responses from neural populations that are spatially intermingled within individual fMRI voxels. Specifically, we factorized the matrix of response magnitudes of each voxel to each stimulus into a set of components, which we hypothesize correspond to distinct neural populations. Each component is described by a response profile across stimuli, and a weight matrix indicating how strongly that component contributes to each voxel’s response (Figure 1). This analysis method enables us to discover the main components that explain neural responses in the ventral visual pathway, potentially including new selectivities not described previously.

**Figure 1:**
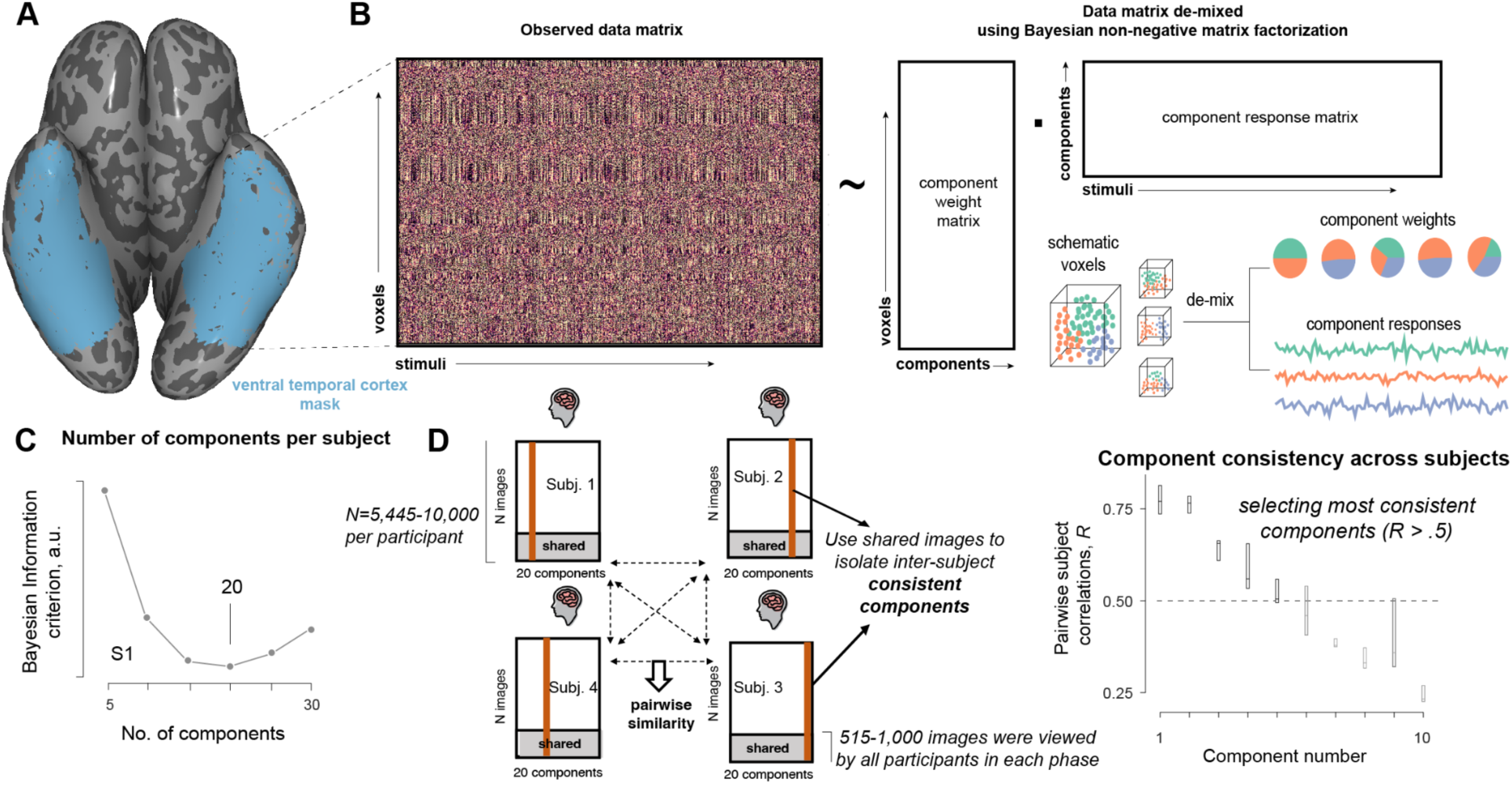
Outline of the data-driven component modeling approach. **A** depicts the large swath of ventral visual cortex included in our analyses for one example subject. **B** illustrates the data-driven voxel decomposition approach. Bayesian non-negative matrix factorization was used to decompose the observed ventral visual stream data matrix of each participant as a product of two lower-dimensional matrices: (i) a response profile matrix that characterizes the response of each component to all 5,445-10,000 stimuli viewed by each participant and (ii) a component by voxel weight matrix that expresses the contribution of each component to each of the ∼6,500-9,600 voxels per participant. **C** shows the Bayesian Information Criterion (BIC) as we vary the number of components in one participant. The optimal number of components was chosen as the minimum BIC. Other subjects had a similar trend, and their data is shown in Figure S1. Components present in all subjects were isolated by measuring pairwise inter-subject correlations of component response profiles, as illustrated in **D** for Phase I participants (right). Gray shaded region shows the proportion of stimuli viewed by all 4 participants in each phase (515 images in Phase 1 and 1,000 in Phase 2). The top 5 components based on this metric all had mean and median pairwise inter-subject consistency > 0.5 (D, right).

## Results

We applied hypothesis-neutral Bayesian nonnegative matrix factorization (NMF)^9^ methods to the NSD^1^ to identify the dominant neural populations in the human ventral visual pathway. Importantly, the algorithm does not have any information about the images or the spatial location of voxels. Instead, it infers the response profiles and anatomical distribution of distinctive neural populations solely from the unlabeled voxel response matrix. This method is thus a powerful way to both validate known selectivities and discover new ones. Our approach is similar to that of Norman-Haignere et al (2015)^8^, except that we use NMF instead of Independent Components Analysis (See Methods for rationale). In Phase I of this project, we analyzed data for four of the eight available NSD participants and presented this work at the Vision Sciences Society meeting^10^. We then pre-registered our analyses on the Open Science Framework (https://osf.io/n47qf) and confirmed our hypotheses on the four held-out NSD participants (Phase 2). We report results for both groups analyzed separately as well as for each individual participant in Supplemental Information.

Our general procedure is illustrated in Figure 1. We first applied the NMF algorithm on each subject’s data separately (Figure 1A, B) to identify subject-specific components (Phase 1 subjects viewed 10,000 images, Phase 2 subjects viewed 5445 images). Bayesian Information Criterion (BIC) applied to Bayesian NMF yielded ∼20 components in each subject (Figure 1C and S1). Next, we used overlapping images viewed by all subjects (in Phase 1 and 2 separately) to identify and rank the consistent components across subjects using a pairwise inter-subject consistency metric. This method identified 5 consistent components across subjects with median pairwise consistency > 0.5. The 5 components derived from Phase 1 participants collectively accounted for ∼50% of the replicable variance in the reliable voxels in Phase 2 subjects (reliability threshold > 0.3, 46% ventral stream voxels), and were highly correlated with the top five components identified independently in Phase 2 subjects (Figure S2), confirming reproducibility of this 5-component structure.

### 1. Characterizing the Function of the Top Components

We first qualitatively examined the response profiles of the top 5 components. For each component, we sorted stimuli by their response magnitude in this component and inspected the top 25 images of each component for each participant. These images (top 4 shown in Figure 2 and all the top 25 shown for each participant in Supplementary Movie 1) revealed a distinctive and familiar selectivity pattern for four of the top five components (Figure 2). The images that produced the highest responses in Components 1, 2, 4, and 5 were, respectively, scenes, faces, text (including words and symbol strings), and bodies (either full bodies or body parts). To validate this apparent preferred category of each component, we collected ratings for each of these preferred categories in a behavioral experiment where participants were asked to rate the salience of each of these categories in each of the images viewed by all NSD participants (see Methods). Salience ratings for the scenes, faces, text, and bodies were strongly correlated with the response of Components 1, 2, 4, and 5 (respectively) across images. These findings are consistent with a large prior literature on selectivities for these categories in the ventral visual pathway^11^ (and their anatomical location, discussed below), so we considered these results as positive controls on our method, and did not interrogate these response profiles further.

**Figure 2:**
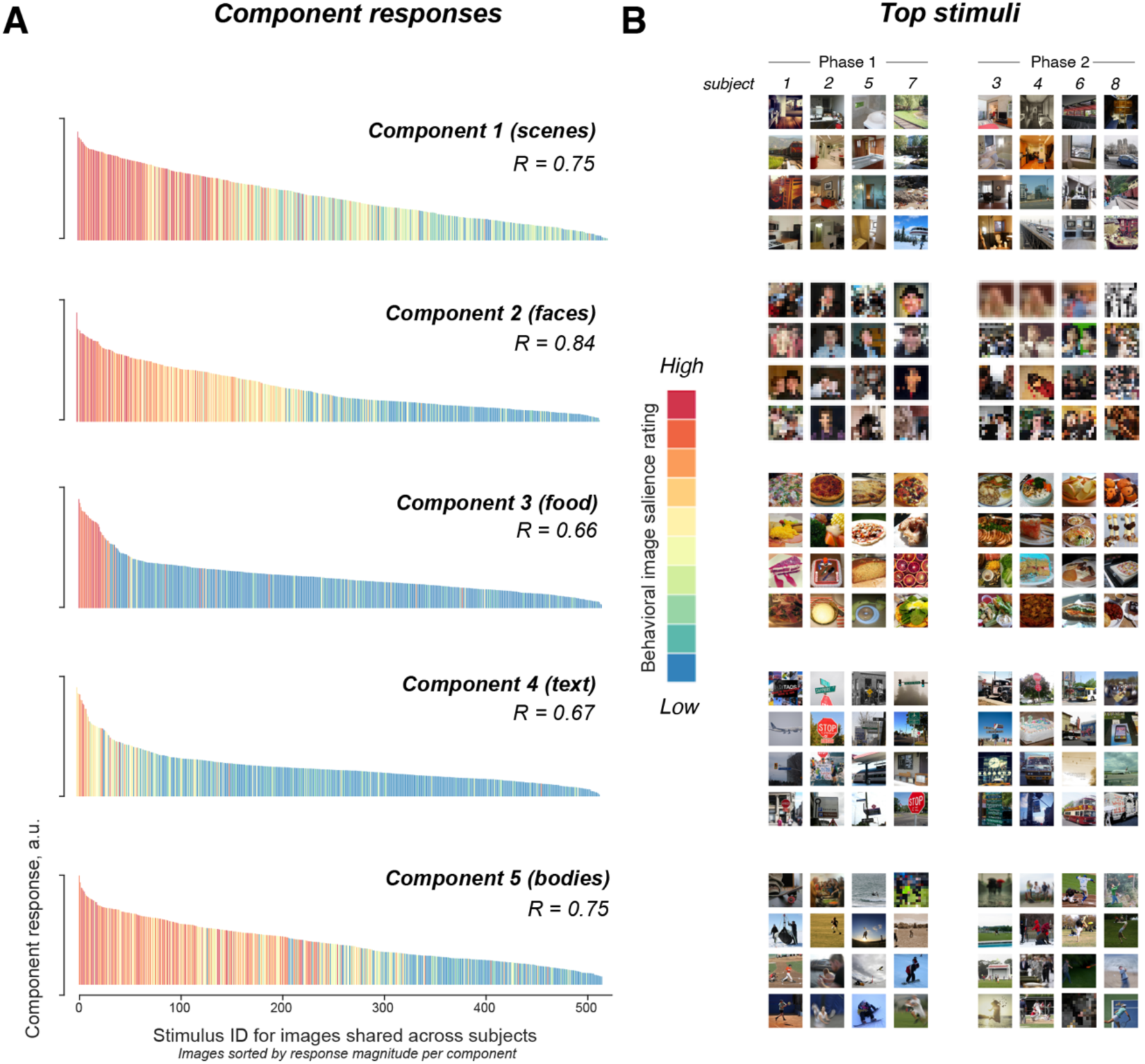
Response profile and preferred stimuli for the top five components. **A.** Response profile for each of the top 5 components (with highest inter-subject consistency) across the 515 images seen by all participants. These components were derived separately within each of the 8 participants individually (see Figure S3 for corresponding data from each participant), but are shown here averaged across all 8 participants. Each bar is an image, and the colors indicate the behavioral salience rating for the preferred category (e.g., the salience of faces for Component 2). **B.** Top 4 images producing the strongest response in each component in each Phase 1 (left) and Phase 2 (right) participant. See Supplementary Movie 1 for the top 25 images for each component in each subject.

### 2. A Novel Component Selectively Responsive to Food

Component 3, however, was unexpected. This component, the third most consistent component across participants in the separate analyses of both Phase 1 and Phase 2 participants, appeared to respond in a highly selective fashion to images of food. This food selectivity is evident both in the correlation of the component’s response profile with rated salience of food (Fig 2A) and in the images that produced the highest response in individual subjects (Fig. 2B). Although most of the top-ranked images are of prepared food (e.g., a slice of pizza), unprepared food (e.g., a broccoli, carrot, banana, etc.) also produced strong responses in this component (see Supplementary Movie 1). But inspection of those top images also suggests several potential alternative accounts for this component’s responses. For example, the top images for this component also seem to share certain low-level and mid-level visual features, including warmer and more saturated colors, higher curvature, and a complex spatial structure with rich texture. To address these potential alternative accounts of food selectivity, we first estimated several image-computable metrics of color, curvature, and texture (see Methods). The component response was more strongly correlated to the behavioral salience ratings for food than any of these other visual feature metrics (Figure 3A). However, some of the visual properties were also significantly correlated with the component response, particularly the object-color probability metric^12^ (the probability of a hue being a natural object, which reflects the warm-cool color continuum).

**Figure 3.**
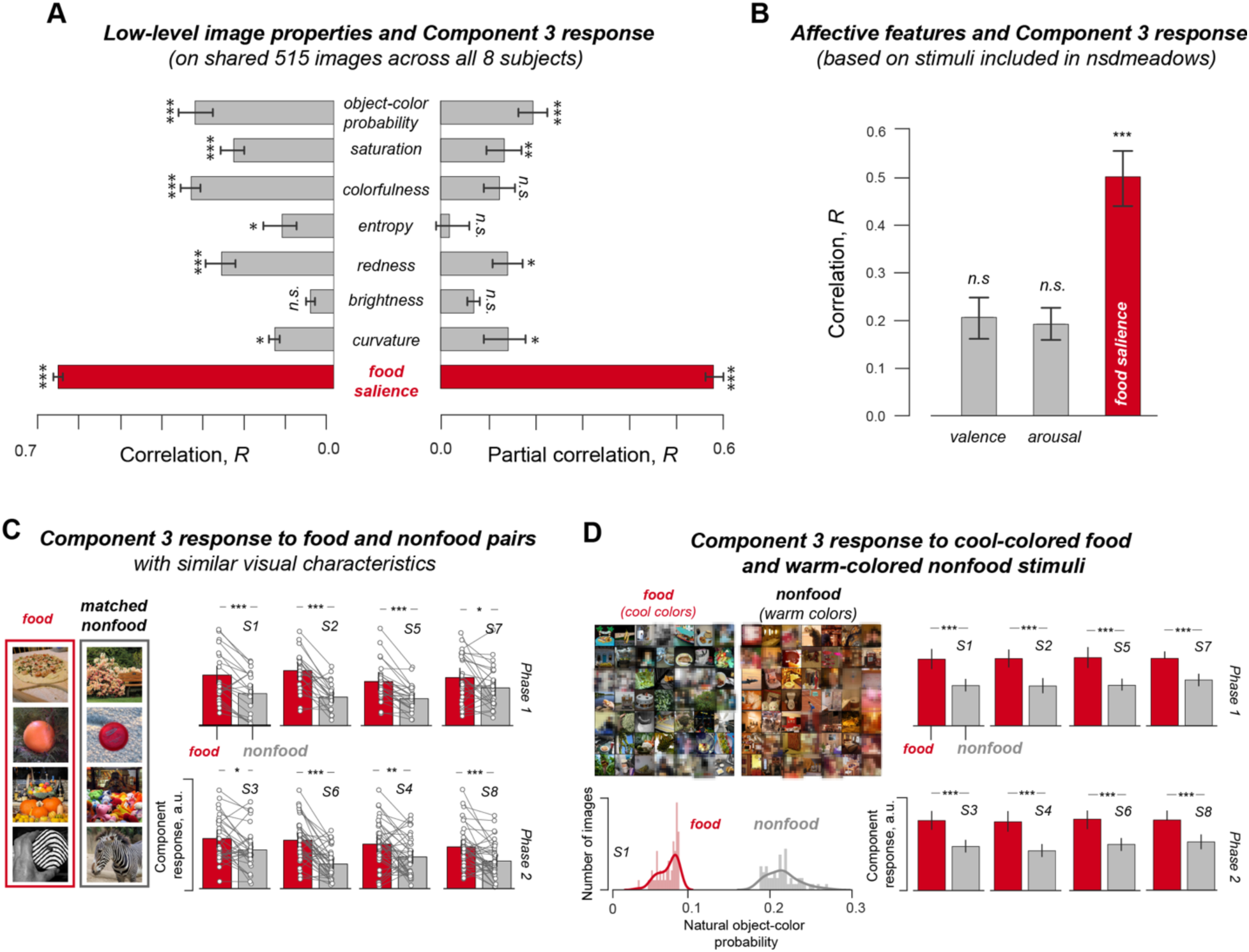
Alternate accounts do not explain the food selectivity of Component 3. **A.** Left: The correlation across stimuli between the magnitude of the Component 3 response averaged across the 8 participants and various image-computable feature dimensions and rated food salience (see Methods). Right: Same as A, but now with food salience partialled out for the image-computable measures, and with all other measures partialled out for the food salience measure. **B.** The correlation between the magnitude of the Component 3 response and valence and arousal ratings across the subset of 100 stimuli for which these ratings were available in the original NSD study (Allen et al 2021). **C.** Responses of Component 3 in each participant to food and nonfood stimuli selected in pairs of images (one food and one nonfood) that produce similar activations in the last convolutional layer (‘*conv5*’) of an AlexNet architecture pre-trained on ImageNet. **D.** Response of Component 3 in each participant to sets of stimuli chosen such that the food images were very low, and the nonfood images were very high on the object-color probability measure (Rosenthal et al., 2018^12^). See also Figure S5.

But how much unique variance do each of these variables explain of the component’s responses? Figure 3A (right) demonstrates that the rated salience of food remains highly correlated with the response profile of Component 3 (*R* = 0.58, *P*=6e-47), even after partialling out all the visual features that appear to be most confounded with food. Some of these visual properties on their own account for a significant though much smaller part (e.g. for object-color probability, R=0.16, p=8e-6) of the Component’s response once food salience is partialed out, although the unique variance explained by food salience ratings is significantly greater than the variance explained by any of these visual properties with the effect of food salience removed (all ps<0.00001, Figure S5). Figure 3B further shows that the response of Component 3 was not significantly correlated with behavioral ratings of either valence or arousal provided for a subset of the stimuli in the original NSD study. Further, the food selectivity of this component persists strikingly across the large set of 5,445-10,000 images viewed by each participant (Figure S4).

These results indicate that the visual features most obviously related to Component 3 cannot alone explain the response of Component 3. However, it remains possible that this component selectively responds to a conjunction of multiple lower-level features (like reddish, round objects). We therefore performed three further analyses. First, we identified pairs of images (one food and one nonfood) from the 5,445-10,000 image set viewed by each participant that produce similar activations in the last convolutional layer (‘*conv5*’) of a pretrained AlexNet (see Methods). These food and nonfood pairs are visually very similar, with matching features in similar spatial locations (see Figure 3C). Yet food images still produced a significantly higher response than their matched nonfood images in each participant (paired t-tests all *P*s<0.01). Second, we identified food images that ranked low on an object-color probability measure (Rosenthal et al. 2018^12^, i.e. ‘cool’ colored food) and nonfood images that ranked high on the same scale (i.e. ’warm’ nonfood images). The component response remained significantly higher to the food images than the nonfood images in every subject (all *P*s<0.001, Figure 3D), suggesting that the component’s food selectivity overrides any sensitivity to object-color probability (see also Figure S5). Third, we selected subsets of food and nonfood stimuli that maximally span the embedding space of different layers of an ImageNet-trained AlexNet model^13^, such that the sampled images within each set are substantially dissimilar among themselves, and the selected subsets are diverse on perceptually relevant image properties (Figure S6; described further in the Methods). As a result, a linear classifier trained to discriminate between these food and nonfood images using the CNN features of the corresponding layer performs at chance (never exceeding 53%). And yet the food images still produce a significantly higher response than the nonfood images in Component 3, even across these highly heterogeneous food and nonfood subsets (Figure S6), showing that the food preference holds broadly and is not limited to specific kinds of food images.

Our analyses thus far focused only on the stimuli included in the NSD. Although the NSD includes a large number of images (N = 56,720 across 8 subjects with full repetitions each), they span a small subset of the space of all possible images. To address this limitation, we built a deep neural network (DNN)-based encoding model to predict the response of Component 3 (see Ratan Murty et al., 2021^14^). Our CLIP-ResNet50^15^ based encoding model was highly accurate at predicting the response to images not encountered in the model training procedure (correlation between the cross-validated predicted and observed responses = 0.83, *P* < 0.00001, Figure 4A and Figure S7). Because this DNN-based model is image-computable and highly accurate, it enables us to test the predicted response of the Component 3 on images beyond those included in the NSD. Would the food selectivity of the component hold even when tested on a much larger battery of stimuli? To find out, we obtained predictions for the Component 3 response to all 1.2 million stimuli from the ImageNet dataset^16^. All the top 1000 stimuli predicted to activate this component (from ∼1.2 million possible images) contained food (see Figure 4B, top for the top 50 images and Supplemental Figure S8 for the top 200 images). This high-throughput screening procedure on DNNs validates the observed food selectivity of Component 3.

**Figure 4.**
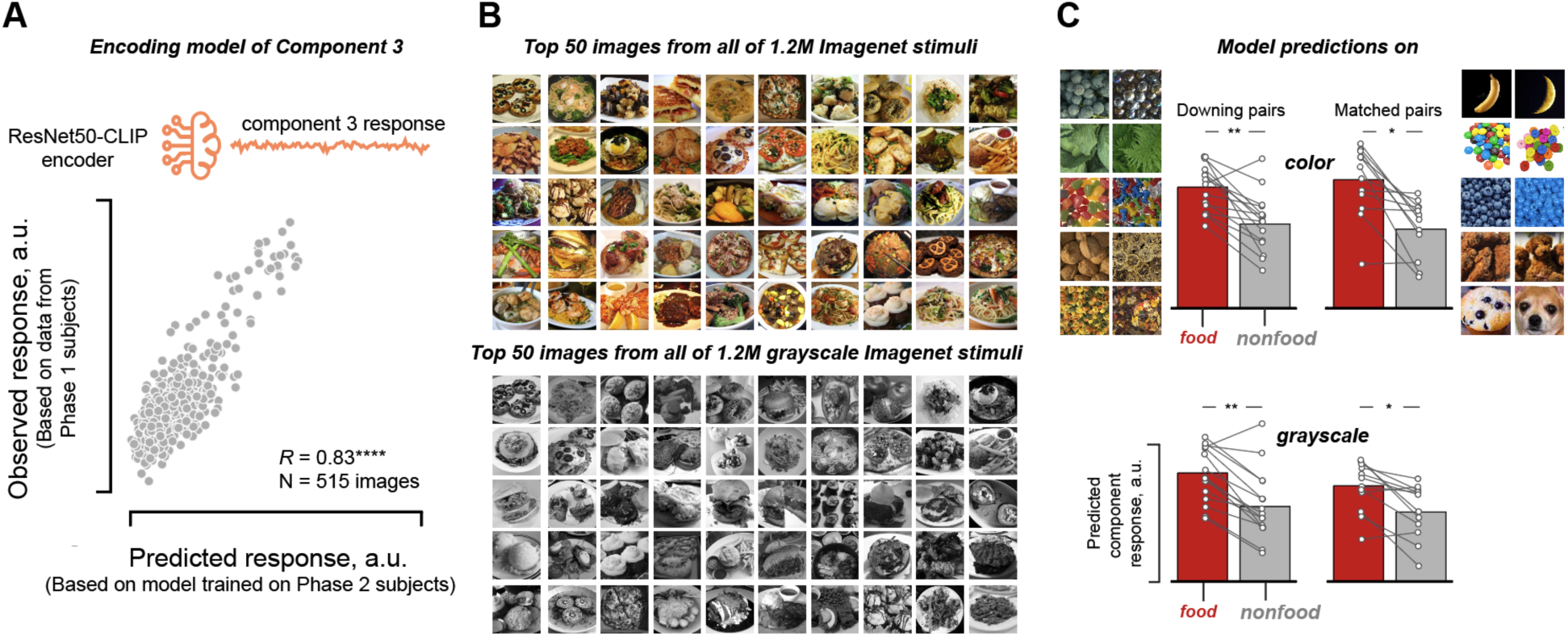
A DNN-based encoding model of Component 3 response enables tests on images beyond those in the NSD. **A.** We used a ResNet50-CLIP encoder to predict the Component 3 response. The x-axis shows the predicted component response (based on the model trained on Phase 2 subjects) and the y-axis shows the observed Component 3 response (from Phase 1 subjects). Each dot is an image (total N = 515 shared images across all 8 subjects) that the model did not encounter in the model fitting procedure (cross-validated on both images and subjects) **B.** Top 50 stimuli predicted by the encoding model to have the highest response across all 1.2M colored (top) and grayscale (bottom) images from the ImageNet dataset. All images are of food. **C.** Model prediction on colored (top) and grayscale (bottom) versions of the Downing pairs and our Matched pairs of food and nonfood images. The bars indicate the mean response, and each connected line indicates a particular matched food-nonfood pair.

Downing and Kanwisher (1999)^17^ had previously tested and rejected the food selectivity hypothesis when they failed to find higher responses to food textures compared to visually similar nonfood textures. When these stimuli were tested on the DNN-based model of the Component 3, it predicted a significantly higher response to food than the nonfood matched textures (Figure 4C (top left), paired t-test t(14) = 5.21, *P* = 1.53 x 10-5). Next, we handpicked a new set of food and nonfood images that look very similar (examples shown in Figure 4C, right). Here too, our computational model predicted a significantly higher response to food than matching nonfood images (Figure 4C (top right), paired t-test t(11) = 3.34, *P* = 0.003). Would food selectivity hold even when tested on grayscale images? We tested this in 3 different ways. First, we obtained predictions for the entire 1.2 million ImageNet stimuli again, but this time on greyscale versions of the same images. The correlation between the predicted response to the color versus grayscale version of each image was very high (N = 1,281,167 images, Pearson *R* = 0.98, *P*< 0.00001). Critically, the top 1000 images predicted to have the highest response even from the grayscale set were all food (see Figure 4B, bottom for the top 50 images). Second, our computational model predicted a significantly higher response to food than the nonfood matched textures from the black and white versions of the Downing image pairs (Figure 4C, bottom left, paired t-test t(14) = 2.93 *P* = 0.007) . Third, our computational model also predicted a significantly higher response to black-and-white food than nonfood in our handpicked matched images (Figure 4C, bottom right, paired t-test t(11) = 2.44, *P* = 0.023) . Together, these computational modeling results complement our previous analyses by confirming the observed food selectivity of Component 3 across a larger number of images, for stringent control images, and by showing that the selectivity of this component for food over nonfood persists even for grayscale images.

### 3. The Observed Components are Not Artifacts of the Stimulus Set Composition

Might the observed components reflect the composition of the stimulus set, rather than a property of the brain itself? Of course, experiments cannot reveal selectivities for stimulus classes that are not included in the stimulus set, and there is likely to be some effect of the relative proportion of different stimulus types in the set. However, it seems unlikely that the category selectivities found (for faces, places, bodies, text, and food) reflect an over representation of these categories in the stimulus set compared to human experience, given that most humans spend at least an hour per day engaged in activities where these visual stimuli feature prominently^18^. In addition, several further analyses show that the food-selectivity of Component 3 is not an artifact of the composition of the stimuli. First, the food-selective component emerges separately in each of the eight participants despite the fact that they saw mostly different stimuli, and it can also be identified in the completely separate BOLD5000 dataset^19, 20^ (Figure 5). And conversely, no food-selective component is found when the same analyses are applied to responses to the same images in retinotopic cortex, dorsal and lateral visual streams or in early layers of a CNN. Thus, the use of the NSD stimulus set on its own is neither necessary nor sufficient to find food selectivity.

**Figure 5:**
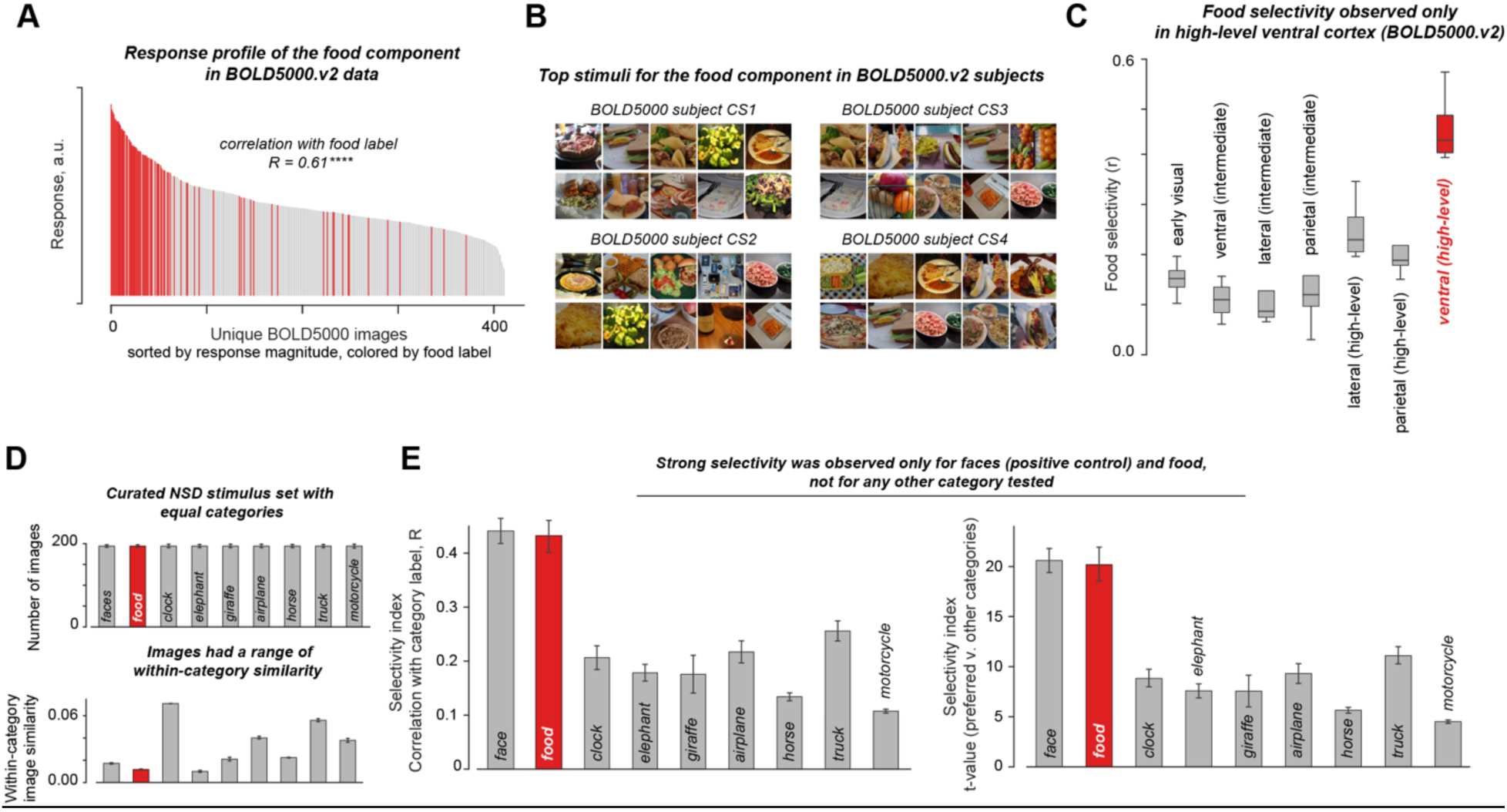
Observed food component is not an artifact of the NSD stimulus composition. We first tested the dependence of stimulus composition, if any, on the completely independent BOLD5000.v2 fMRI dataset. **A.** Response profile for the food component identified in the BOLD5000 data (from images shared with NSD) on images unique to BOLD5000.v2. The x-axis shows the stimuli, and the y-axis shows the inferred response magnitude for the Component 3 for images unique to BOLD5000. The bars in red are images that were labeled as food and the bars in gray are images labeled nonfood in the (imperfect) annotations provided with MS-COCO. **B.** Top 10 images for each of the four subjects in the BOLD5000v2 dataset. **C.** Boxplots showing the food selectivity distribution across BOLD5000 subjects (y-axis) for the components inferred from different cortical regions (x-axis). Food selectivity was observed only in ventral visual cortex, not in other regions. **D.** Next, we performed the NMF decomposition on a curated subset of the NSD with 9 stimulus categories, each with an equal number of images within each subject (top) with high within-category visual similarity for the nonfood categories (bottom). **E.** NMF decomposition on this curated stimulus set revealed components with strong selectivity only for faces (positive control) and food, not for any other category. The y-axis shows the highest selectivity obtained for each of the 9 categories (amongst all components) based on category labels from MS-COCO using two different metrics (left, correlation with a binary category label vector indicating whether the category was present/absent in the image; and right, t-value comparing the mean response to stimuli from that category versus all other stimuli). See Methods for details.

Finally, to test whether any stimulus category that represents a sizable proportion of the stimuli will result in a component selectively responsive to that category, we performed one further analysis (Figure 5D,E). First, we selected a subset of the stimuli that contained equal numbers of exemplars in each of 9 categories including faces (as a positive control), food, and 7 other perceptually homogeneous categories (airplanes, clocks, horses, elephants, giraffes, trucks, motorcycles). Repeating the NMF analysis on these data revealed one component selectively responsive to faces and another to food, and no components as highly selectively responsive to any of the other categories, even though the food images were drawn to be one of the least homogenous in this set. Thus, ample representation of a category, even a perceptually homogeneous category like airplanes or clocks, in the stimulus set is not sufficient for a component selective to that category to emerge, and cannot account for the food-selectivity of Component 3.

### 4. Demixing reveals stronger selectivity for components than voxels

Why was food selectivity not observed before, particularly in previous hypothesis-driven investigations^17, 21^? We speculated that the spatial overlap of food-selective neural populations with other selectivities dilutes food selectivity in individual voxels which our demixing procedure is able to uncover. We tested this idea by measuring the selectivity of the demixed Component 3 and of the average response across the top 1% voxels with the highest weight on Component 3. Food selectivity was significantly higher for the demixed Component 3 (mean *R* across subjects=0.53, P<0.00001) than in the top voxels (mean *R*=0.41, P<0.00001) in each participant (t(8)=10.7, p<0.0001, Figure 6). For comparison, face selectivity of Component 2 was as strong in the top voxels as within the inferred component response (t(8)=0.85, p=0.42, Figure 6). These results indicate that the neural populations selective for food are likely more mixed with other neural populations within voxels than the face-selective neural populations, explaining why strong food selectivity has not been found previously with standard analysis methods.

**Figure 6:**
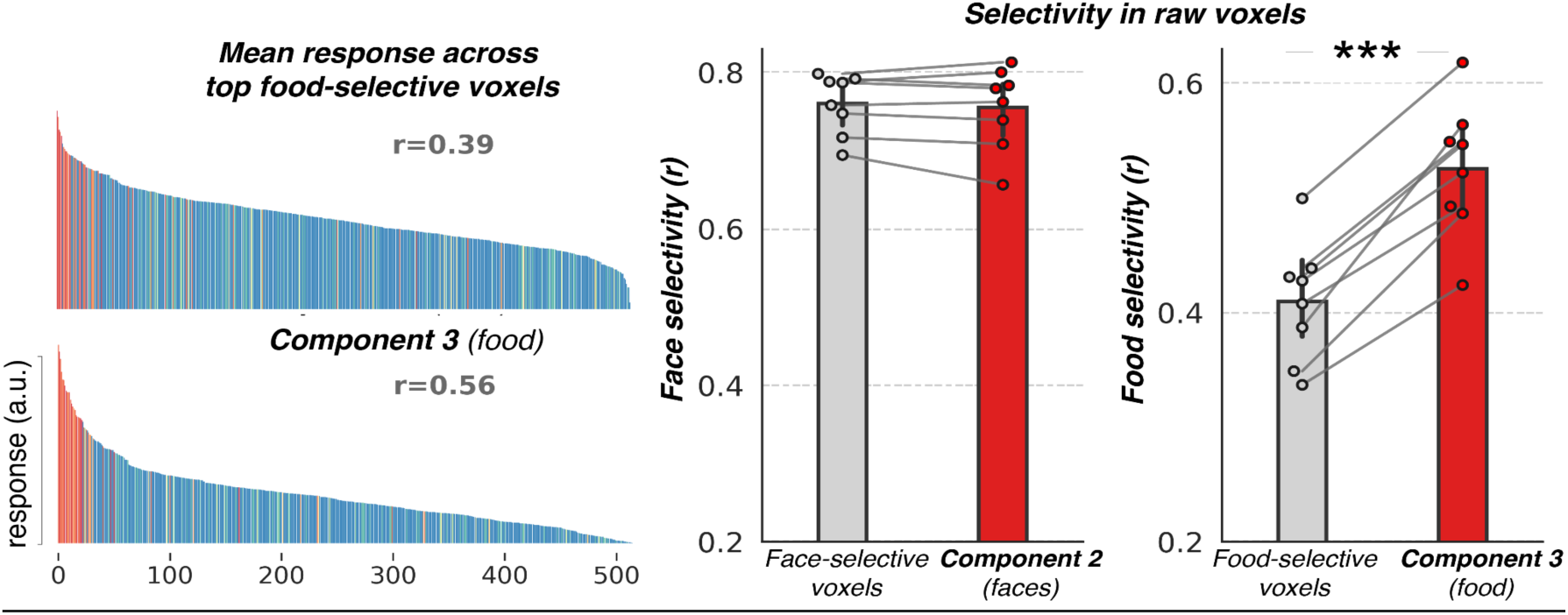
Stronger Selectivity for Components than Average Response across Voxels. Left: Average response profile of the top 1% voxels with the highest weights on Component 3 (top) and the response profile of Component 3 (bottom) for one subject. Food-selectivity of the respective response profiles is reported at the top of each subplot. Right: Face selectivity of the mean responses across top 1% voxels with highest weights on Component 2 (the face-selective component) and the corresponding selectivity of Component 2. Rightmost plot shows the food selectivity of the mean response across top 1% voxels with highest weights on Component 3 (the food-selective component) and the corresponding selectivity of Component 3. Each dot in the swarm plot is an individual subject. Here, selectivity is computed as the correlation between responses and the salience ratings for the preferred category over the 515 stimuli shared across all 8 participants.

### 5. Anatomical Distribution of Components

We next characterized the anatomical distribution of each component by projecting its voxel weights back into anatomical coordinates within each participant individually. For known selectivities, the component anatomies exhibited clear agreement with the corresponding regions identified with an independent functional localizer: the face component produced highest voxel weights in the fusiform face area (FFA) and other face-selective sites like aTL-faces (anterior temporal lobe faces) and mTL-faces (mid temporal lobe faces), the word component was concentrated within the visual word form area (VWFA) and the bodies component in parts of FFA and the extrastriate body area (EBA), as shown qualitatively in Figure 7 and Supplementary Movie 2. Quantitatively, the voxel weight maps demonstrated high correlations with the t-statistics pertaining to the relevant domain from the functional localizer experiment (Figure S9).

**Figure 7:**
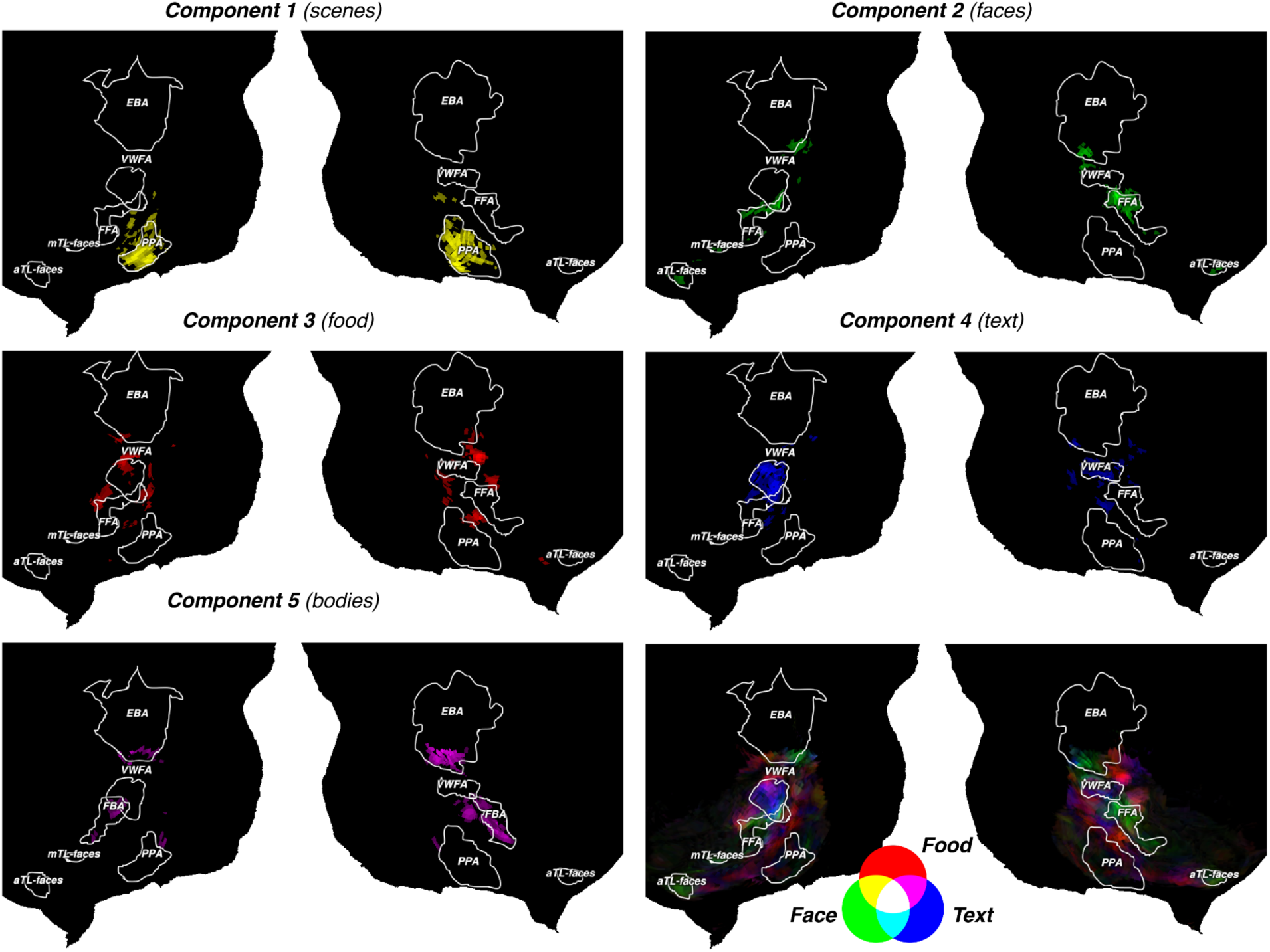
Anatomical locations of highest weights for each component. Top 5% voxels with the highest weights on each component are visualized on cortical flatmaps for one subject. Established regions of interest, defined from the functional localizer scans by computing the contrast of preferred versus all other stimuli, are shown in white outlines (t-value > 2.5). Lower right: Voxel weight maps of face, food and text-selective components are visualized together on the RGB colormap to show component overlap for the same subject. Similar maps for the remaining subjects are shown in Supplementary Movie 2.

The weight maps for the novel food-selective component appeared patchy across the cortex, with considerable variability across participants. To quantify this impression (Figure S10), and compare it to other components, we first registered each participant’s voxel weight map to a common MNI space, and then measured the correlation of the weight map for each participant with the average weight map across the 7 other participants (averaged across 8 folds). This analysis showed the highest inter-subject correlation of the weight maps for the Component 1 (scenes), followed by Components 2 and 4 (faces and text, respectively) and then Components 3 (food) and 5 (bodies). We next quantified the spatial distribution of the voxel weights for each component using a sparseness measure based on the relationship between L1 and L2 norms^22^. Component 3 was less sparse than the others, except for Component 5. Kurtosis and skewness of the voxel weight distributions of each component show that all components have voxel weights that are positively skewed and kurtotic, relative to a Gaussian, indicating a peakier, heavy-tailed distribution skewed towards higher values. A measure of lateralization of the weight maps showed that the face component was right lateralized, and the text component left lateralized, (as expected). The food component trended toward left lateralization, although the lateralization effect was found to be non-significant (one-sample one-tailed t-test: t(8)=1.76, p=0.06). Taken together, these analyses indicate that the inter-subject variability, the sparseness, and degree of the lateralization of the food component is within the range of the other components, but at the lower end of that range.

Given the small but significant correlation across stimuli of the Component 3 response with various measures of color information, even after food salience was factored out, we next asked how similar the anatomical distribution of this component was to the distribution of color responsiveness across the cortex^23, 24^. Specifically, we measured the correlation between the saturation of nonfood stimuli with voxel responses in the VVC to those stimuli, and then compared the resulting correlation map with the voxel weights for Component 3, separately within each participant. We find (Figure S11) that the saturation-responsiveness map is indeed correlated with the Component 3 weight map in every subject (mean ∼0.4), and more so than for any other component (p < 0.01 for all 4 comparisons using a paired t-test). Thus, the anatomical distribution of Component 3 and color responses are correlated with each other across the cortex.

## Discussion

We applied data-driven analyses to a very large data set of fMRI responses to thousands of natural images and found that the dominant neural response profiles in the ventral visual pathway include selective responses to faces, scenes, bodies, text, and food. Although the first four of these selective responses have been reported in many previous studies, what is novel in the current study is their emergence unbidden from a hypothesis-neutral analysis of a dataset that was not designed to test for or reveal them. The fact that these four previously reported selectivites emerged separately within each of the 8 individual participants in this study (each of whom saw mostly non overlapping images), shows that they reflect not just the idiosyncratic whims of the scientists who chose to test these hypotheses in the past, but the actual dominant features of the neural response in the ventral visual pathway. But our most novel result is the discovery of a new neural response that has not been reported previously for the ventral visual pathway that is highly selective to images of food. Taken together, these results give a more comprehensive and data-driven characterization of the dominant neural response profiles of the ventral visual pathway, describe a new neural selectivity for visual food images, and provide new clues into why we have the neural selectivities we do.

Because our finding of neural selectivity for food was unexpected, we embarked on an extensive series of control experiments to test alternatives to this hypothesis. We found that although the magnitude of response of Component 3 was correlated with the presence of visual features such as color saturation, warm colors, curved shapes, and texture properties, the only factor that remained highly correlated with the food component response when other factors were partialled out was the salience of food in the image (Figure 3A). Second, when we pitted the presence of warm colors against food salience as accounts of the response of this component, we found that food trumped color: cool-colored food produced a higher response in this component than warm-colored nonfood. Third, the food selectivity of this component persisted even for computationally-matched stimulus pairs (one food, one nonfood) that elicit similar activation patterns in deep layers of a pre-trained CNN. Fourth, we built a CNN-based model for Component 3^14^, which accurately predicted responses of this component to held-out stimuli, and we used this model to turbocharge our search for counterevidence of the food selectivity of this component. We ran all 1.2 million images from ImageNet through this model and looked at the top 1,000 predicted to produce the highest response. They were all food. Fifth, we constructed pairs of visually similar food and nonfood images by hand (e.g., a yellow crescent moon and a banana), and ran these images through our predictive model of Component 3. Again, predicted responses were higher to the food images than to their paired visually similar nonfood images. Sixth, an analysis of responses to a subset of nine stimulus categories, including an equal number of stimuli in each category, and importantly, only including food images that are maximally distinct from each other, revealed components selectively responsive to faces and food, but no components responsive to the other 7 categories, showing that the observed selectivity for food is not an artifact of its over-representation in the stimulus set or of the homogeneity of specific kinds of food in the dataset. Taken together, these analyses argue that Component 3 is selectively responsive not to any particular visual features, but to food per se. We have therefore labeled this inferred neural population the Ventral Food Component. Confirming this finding, two preprints based on the same NSD fMRI data finding voxelwise selectivity for food and their overlap with color-biased regions appeared recently^25, 26^. Our study using hypothesis-neutral methods further shows that food selectivity is a dominant feature of ventral visual cortex, that food selectivity is strong when revealed by demixing methods, and that food selectivity overrides low and mid-level features, including color.

While a distinctive response to visual images of food has been described in taste-sensitive regions of the insular cortex^27^, a robust and highly food-selective response in the visual cortex has never been observed before. Why has the food selective component not been reported previously, despite past efforts to look for it^21, 28, 29^? The likely account is that prior studies primarily analyzed raw voxel responses, which do not reveal strong selectivity for food, because the food component is spatially intermingled with other neural populations within voxels. In contrast, our voxel decomposition method demixes these neural responses, revealing the strong selectivity of the food component alone. Indeed, when we obtained the visually matched pairs of food and nonfood textures that had produced similar responses in Downing and Kanwisher’s (1999) study^17^, leading them to argue against food selectivity, we found that our predictive model for Component 3 produced a higher response to food than nonfood. A similar pattern was observed previously with our voxel decomposition analysis of responses in auditory cortex^8^, where we found only weak selectivity for music in raw voxels, but strong selectivity for music in the inferred music component, which was later validated by clear music selectivity in the responses of individual intracranial electrodes^30^.

A notable property of Component 3 is that even though its selectivity for food cannot be explained by responsiveness to color properties alone, the two are clearly linked. The response profile of Component 3 has a much smaller but still significant correlation with color metrics even after food selectivity has been partialled out, and its anatomical distribution across the cortex is correlated with the anatomical distribution of responsiveness to color information (see Figure S11). Why might the apparently same neural population be responsive to both food and color information, even when each is unconfounded from the other? Many have noted the importance of color for the detection, evaluation, and choice of food^31, 32^. Neuropsychological studies of patients with cortical color blindness (achromatopsia) have noted particular difficulties in discriminating food. Pallis (1955)^33^ quotes an achromatopsic patient saying, “I have difficulty in recognizing certain kinds of food on my plate. I can tell peas and bananas by their size and shape. An omelette, however, looks like a piece of meat.” Further, behavioral studies have shown that adults, preschool children^34^, and monkeys^35^ use color more than shape when generalizing across food categories, but the opposite when generalizing across nonfood categories. Indeed, Santos et al. (2001)^35^ argued that the use of color over shape only in food learning suggests the existence of a domain-specific mechanism for visual food choice. Studies in typical adults also reinforce the deep link between color and food perception, finding for example that images of food (but not nonfood) are rated as having higher arousal if they contain red, and lower arousal if they contain green^31^. The authors of that study speculate that red color was indicative of the caloric and nutritional value of food for our evolutionary ancestors but is much less so today (in prepared foods and where food dyes are used), and hence reveals the evolutionary basis of the connection between color in general and red in particular in food preference. Of course, food preferences are famously culture-specific and learned^36^, and infants’ food choice is primarily learned from other people. But color may still play a role in domain-specific learning about food, and in bootstrapping the development of a cortical circuit for visual food discrimination. One hypothesis is that the color bias in food choice may arise relatively early in development (though not apparently in infancy^34^), with the cortical locus of the Ventral Food Component accordingly arising in regions already biased for warm colors^12^, but that the particular visual food stimuli that activate this system are most likely learned through individual experience (like orthographies in the VWFA).

What does food selectivity tell us about which categories get their own specialized neural machinery in the brain? Because food has been of fundamental importance to humans both throughout their evolution, and in modern daily life^36^, and because food choice often starts with vision, a specialization for food in the visual cortex is consistent with both evolutionary and experiential origins of cortical specializations. On the other hand, food seems more visually heterogeneous than other categories with selective responses in the ventral pathway, an impression confirmed by visual similarity measures based on feature responses in pretrained AlexNet (Figure S12). Nonetheless, food is linked to some visual features, notably color, and indeed we find that Component 3 does show a small but significant color preference even after food salience is partialled out. This finding is reminiscent of other feature biases in category-selective cortex (e.g curvature biases in face selective cortex^37^), and invites the same chicken-and-egg question: Do category selectivities colonize cortical regions with pre-existing relevant feature biases^37^, or are these visual feature preferences simply by-products of category selectivity? Finally, the finding of food selectivity resolves a previous conflict with the hypothesis that category selectivity in visual cortex is determined by the computational requirements of the task^38^. We had proposed this hypothesis in a recent study^38^, based on our finding that convolutional neural networks trained on both face discrimination and object classification spontaneously segregated themselves into separate systems for face and object recognition. But that study also found spontaneous segregation for food in a network trained on both food and object classification, a finding that seemed then not to fit the brain, but that now does. Thus, the novel selectivity for food reinforces the computational hypothesis that task constraints play a role in determining which categories are processed with their own specialized neural machinery.

What computational advantages might a selective response to food confer? Any form of selectivity in a neural population inherently implies a sparse code, as it suggests that the neural population responds strongly to only a specific subset of all possible stimuli. Such sparse neural codes have long been argued to make information explicit and easier to read out^39–41^, and to support faster learning^42–44^. Reducing metabolic costs would also favor sparse codes for stimuli that are most frequently encountered in our environment. Of course, we cannot have specialized neural codes for all possible classes of stimuli, so it would be sensible to allocate such specialized systems to a relatively small number of the most important object classes - like food.

A final note is that our analysis did not find evidence for selective neural responses to several visual features and categories for which ventral visual pathway specializations have been proposed in the past, including animals^45^ (which are well represented in the stimulus set) and stubby-shaped and spikey-shaped objects^46^. We also did not see evidence for previously-proposed selectivities for small inanimate objects^45, 47^ or tools^48, 49^, although these selectivities may be located more on the lateral than ventral surface of the brain, outside the search window used here. Of course, there are many reasons why selectivities that exist in the brain might not be detected using fMRI, but the failure of previous findings from fMRI to emerge from the current analysis raises questions about whether those selectivities might already be better accounted for by the components found here^50^.

In sum, our hypothesis-neutral investigation on the ventral visual pathway reveals neural populations selective for faces, places, bodies, text, and food. The fact that these selectivities emerge from a hypothesis-neutral analysis, across multiple largely non-overlapping sets of images, indicates that they reflect not just the capricious interests of researchers in the past, but rather constitute dominant features of the functional organization of the ventral visual pathway. Further, the novel selectivity for food reported here raises fascinating questions about its developmental origins, connectivity, and behavioral consequences. Another important open question is whether this neural population represents the mere presence of food, or its familiarity, appeal, or caloric or nutritive content. This new finding further shows that selective neural responses in the ventral visual pathway arise not only for perceptually homogeneous categories that may reflect confluences of overlapping visual feature maps^51^, but also categories that are visually quite heterogeneous, especially if exemplars of that category require specialized computations for their discrimination^38^ and engage our frequent and abiding interest.

## Acknowledgements

This work was supported by National Institutes of Health grant DP1HD091947 to Nancy Kanwisher, K99/R00 Pathway to Independence Award from the National Eye Institute of the NIH (grant 1K99EY032603) to NARM, and the Center for Brains, Minds, and Machines (CBMM) funded by NSF STC award CCF-1231216. This work was presented on May 15, 2022, at the Vision Sciences Society (abstract submitted December 2021). We thank Elizabeth Meiczkowski for help obtaining behavioral data and members of the Kanwisher lab for helpful suggestions on the manuscript.

## Author Contributions

NK and MK conceived the study. MK performed the main NMF analyses in the paper and RM built the DNN model and performed analysis on its predictions. NK, MK, and RM wrote the paper.

## Methods

### Natural Scenes Dataset

A detailed description of the Natural Scenes Dataset (NSD; http://naturalscenesdataset.org) is provided elsewhere (Allen et al., Nature Neuroscience, 2021^1^). Briefly, the NSD contains measurements of fMRI responses from 8 participants who each viewed 9,000–10,000 distinct color natural scenes (22,000–30,000 trials) over the course of 30–40 scan sessions. Scanning was conducted at 7T using whole-brain gradient-echo EPI at 1.8-mm resolution and 1.6-s repetition time. Images were taken from the Microsoft Common Objects in Context (COCO) database^52^, square cropped, and presented at a size of 8.4° x 8.4°. A special set of 1,000 images were shared across half the subjects with full repetitions (participant NSD IDs: 1,2,5,7) and a subset of these (515 images) were shared across all 8 participants. The remaining images were mutually exclusive across subjects. Images were presented for 3 s with 1-s gaps in between images. Subjects fixated centrally and performed a long-term continuous recognition task on the images. The fMRI data were pre-processed by performing one temporal interpolation (to correct for slice time differences) and one spatial interpolation (to correct for head motion). A general linear model was then used to estimate single-trial beta weights. Cortical surface reconstructions were generated using FreeSurfer, and both volume- and surface-based versions of the beta weights were created. In this paper, we used the 1.8-mm volume ‘nativesurface’ preparation of the NSD data and version 3 of the NSD single-trial betas (betas_fithrf_GLMdenoise_RR).

#### Additional Data preprocessing

We analyzed responses only to images that were seen three times in the participant in question. We averaged single-trial betas across the 3 repetitions after z-scoring every voxel separately within each scan session to create our voxel responses. This within-scan normalization was performed to account for differences in mean percent signal change (PSC) across scan sessions, which may arise due to incidental variability in global BOLD signals.

We extracted high-level ventral visual stream voxels by using the *streams* atlas provided in the native space of each subject with NSD. This atlas is largely based on fsaverage folding but also accounts for the noise ceiling to ensure that the regions cover reliable voxels. To make the data matrix suitable for NMF so that it contains all positive entries, we perform a baseline shift of voxel responses by subtracting the minimum z-scored response of each voxel (across all stimuli) from its responses to all stimuli.

### A Bayesian Matrix Factorization approach for the analysis of large-scale fMRI recordings

We model the data matrix (voxels x images) as the product of two lower rank matrices. The first matrix (called the response profile matrix henceforth) encodes the response profiles of each component (‘neural populations’) to all images and the second matrix (called the component by voxel weight matrix) specifies the relative contribution of all voxels to each component. We chose NMF for our matrix factorization algorithm for several reasons. First, PCA/ICA based approaches do not yield “signed” components, i.e., negative and positive weights are treated equivalently. Initial pilot analyses of our data using the PCA/ICA approach of Norman-Haignere et al. (2015)^8^ revealed single components with both positive and negative responses and voxel weights which couldn’t be oriented (or flipped) such that the response and voxel weights predominantly have the same sign. Negative response magnitudes are generally not consistent with neural response in the ventral visual pathway, which usually increase after stimulus presentation, and negative voxel weights violate our modeling assumptions about the voxel weight matrix representing the relative anatomical proportions of each component in every voxel. Further, ICA requires the statistical independence of unmixed components and can fail in practice when no linear demixing matrix is found, as can happen when there is significant spatial overlap between distinct neural populations and independence is not achievable. NMF is better equipped to handle spatially overlapping signals in such cases, and has been more effective in other neuroscience domains that rely on demixing of spatially overlapping components^53^. Importantly, NMF makes only minimal and biologically meaningful assumptions about the components by enforcing the basis functions to be nonnegative. These considerations led us to favor an NMF-based approach over other decomposition techniques. Since choosing the number of components is an important problem in NMF, we adopted a Bayesian NMF approach^9^ since it affords a principled way of selecting the number of components based on likelihood (e.g., Bayesian Information criterion). For a comparison of the food selectivity of the NMF-derived food component with the most food-selective PCA component and cluster (given by k-means clustering), see Figure M1.

Mathematically, the Bayesian NMF algorithm models the data matrix ***D*** as,

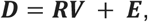

where **D** is the images x voxels data matrix for every participant, **R** is the image x components (N x C) *response profile matrix*, **V** is the components x voxels (C x V) *voxel weight matrix* and **E** is an images x voxels (N x V) *residual matrix*. In the Bayesian approach to NMF, all parameters for (**R, V, E**) are stated in terms of their prior densities. For efficient inference, following Schmidt et al., (2009)^9^, we choose a zero mean normal residual matrix **E** with variance *σ*^2^, and a normal data likelihood,

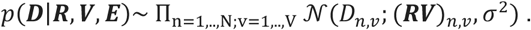

Further we assume that **R** and **V** are independently exponentially distributed with scales *ρ_n,c_* and γ*_c,v_*

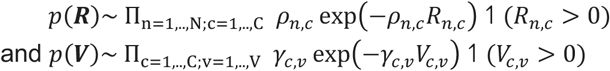

The conditional probabilities of **R** and **V** thus have a rectified Gaussian distribution. Following Schmidt et al., (2009)^9^, the prior for the variance in **E** is assumed to have an inverse gamma distribution, resulting in an inverse-gamma conditional probability. Parameters for (**R, V, E**) are optimized by sequentially drawing samples from these conditional densities using the Bayesian Markov Chain Monte Carlo (MCMC) sampling method derived in Schmidt et al., (2009)^9^.

**Figure M1.**
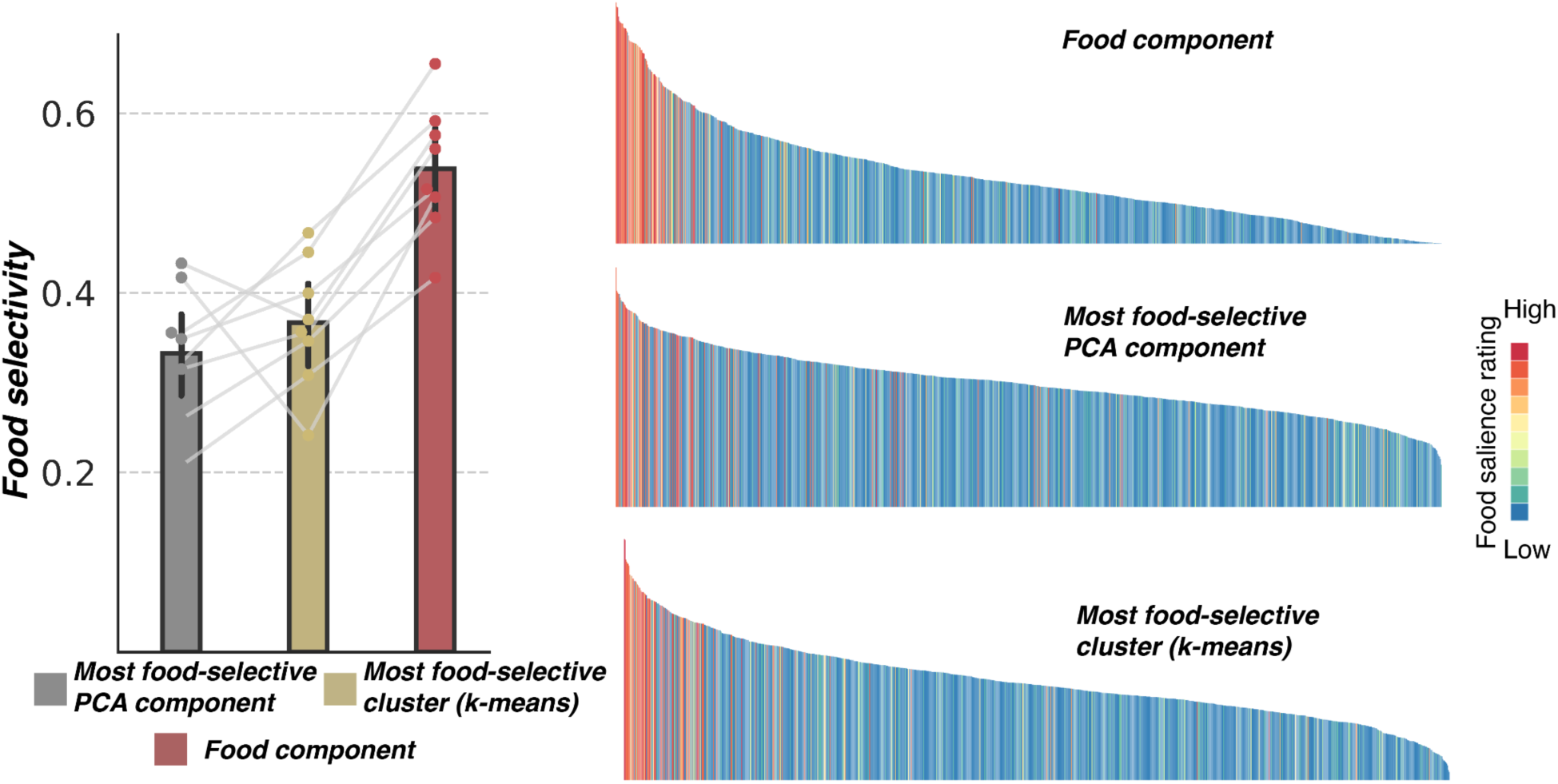
Food selectivity of the NMF-based Component 3 is compared against the selectivity of the PCA component (among top 20 components) and cluster (as partitioned with k-means clustering using k=20) with the highest food-selectivity. Selectivity is computed by correlating the response profile of each component/cluster with the salience ratings for food. Right panel shows the respective response profiles colored by food salience ratings for one representative subject.

### Extracting robust components in individual subjects with a consensus approach

Like standard NMF, Bayesian NMF is also a stochastic algorithm sensitive to initialization and accurate initialization of the estimates is critical. To get robust components, we run this algorithm N= 50 times on the data matrix for each subject to get C=20 components per run. We then perform a consensus NMF procedure inspired by Kotliar et al. (2018)^54^ to aggregate results from different runs of the NMF algorithm into a single stable matrix factorization result. In this procedure, the estimated response profile matrices from each run are concatenated across the component dimension to create an (images x NC) matrix where each column is a component from a single run of the algorithm. We follow the same procedure as described in Kotliar et al. (2018)^54^ to get the consensus response profile matrix (images x C) from this aggregated data matrix. This consensus algorithm first isolates and removes unreliable components by running an outlier detection procedure, enabling us to filter out components that are not replicable across runs. Next, the remaining components over all runs combined are clustered (with C clusters) and the medians of these clusters are returned as the consensus (stable) response profiles of the C components.

The final voxel weight matrix for each subject is then obtained by finding component indices in individual NMF runs that have the highest correlation with each of the C consensus NMF component response profiles. The respective voxel weights for each index are normalized (to sum up to 1) and then averaged across runs. This gives us the consensus voxel weights for each component.

### Extracting components with high inter-subject consistency

The previous analysis yielded 20 components in each individual subject. Since we are interested in discovering the functional organization structure *shared* across individuals, we next analyzed the one-to-one correspondence between these components across subjects. To determine which of the resulting components for each participant are shared across participants, we use the 1,000 images (or 515 images in the case of Phase II participants) that were viewed by all participants. Specifically, we rank-ordered components based on the highest average inter-subject correlation in their response to the shared images. Since there are 4 subjects in each phase of our analysis, we get 6 unique pairwise correlation values for every possible combination of ordered component indices across the 4 subjects (20 x 20 x 20 x 20). The inter-subject correlation measure, called ‘inter-subject consistency’, is computed as the average of these 6 values. We first pick the component indices (i, j, k, l) that yield the highest inter-subject consistency. We then repeat the same procedure on the (19 x 19 x 19 x 19) matrix after removing the indices (i, j, k, l) and repeat this procedure until the inter-subject correlation drops significantly. As shown in Figure 1, this value drops sharply after a handful of components, and we restrict our analysis to the top 5 components which all demonstrate an average inter-subject correlation value of 0.5 or greater.

### Independent replication with held-out subjects data

The hypothesis that the ventral pathway contains a neural population that responds selectively to food was formulated based on analyses of the data from four participants in Phase 1 (NSD participants 1,2,5,7), before hypotheses and analysis methods were registered on OSF, and then tested on the held-out data in Phase 2 (NSD participants 3,4,6,8).

Only the top few images for the 5 most inter-subject consistent components were visually inspected to ascribe semantic categories to components. The same top 5 components (with top images respectively selective for faces, scenes, food, text, and bodies) were obtained in the Phase 2, confirming reproducibility of our findings and allowing us to combine responses across the different phases.

This apparent category selectivity of each of the top 5 components was subsequently rigorously assessed using quantitative measures based on salience ratings, as described below.

### Behavioral experiment

We collected subjective salience ratings from Amazon Mechanical Turk for the 1,000 images viewed by all the Phase I participants. Among these images, 515 images were also viewed by all Phase II participants with three repetitions. In this experiment, participants were asked to rate the salience of each of the 5 categories that seemed to be intuitively represented in the top images for each component, namely, *scenes, faces, bodies, text,* and *food*. Specifically, participants were given the following task: ‘Rate how prominent [category] is within each image’ and were instructed to provide a rating on a scale of 0 to 9. Each participant completed salience ratings for one category over a series of 220 images and we obtained 5 ratings per category (from 5 participants) and averaged the ratings across participants to get an overall measure of the salience of each category for each image. Salience ratings for scenes were odd, presumably because most natural images have some kind of scene context, and participants were unsure what we meant. Therefore, for the scene category only we instead asked two experts in scene-selective cortex who were uninvolved in the study to rate their prediction for how strongly the image would drive the scene-selective cortex.

### Component selectivity analysis

We performed a correlation analysis (Pearson’s r) to quantify the extent of agreement between the responses of each component and the salience ratings of their preferred category (as visualized in the top images) over all 515 images that were viewed by all participants. Component responses were first averaged across all 8 subjects before correlation computation (Figure 2). We further also computed these correlations at the single-subject level using all stimuli for which the salience ratings were available (1,000 for Phase I participants and 515 for Phase II participants), as shown in Supplementary Figure S3.

### Control analyses on the novel component

I. Image-computable properties: We computed the following image-level properties to assess their respective impact on driving food component responses (our alternative accounts for food selectivity). These properties include:

**Color metrics:**

a. Saturation: Mean saturation of every image is computed after transforming the image from RGB to HSV space
b. Brightness: Brightness is computed as the mean value across the ‘V’ channel after transforming the image from RGB to HSV color space.
c. Colorfulness: This metric is included to capture the *perception* of colorfulness. We compute the colorfulness metric for every image based on the opponent color space representation discussed in Hasler and Suesstrunk (2003)^55^.
d. Hues (Redness): The histogram of the hue channel is computed after binning the hue values across all spatial locations in the image into 8 equally spaced radial bins. The top hue (among the 8 bins) that had the highest correlations with the food component response roughly corresponded to red hues. We thus included the hue values in this bin in the subsequent partial correlation analysis while assessing the unique contribution of each metric in explaining the food component response.
e. A color representational axis defined in Rosenthal et al. (2018)^12^, called ‘Object-color probability’ is computed as the probability of a given hue being a natural object in an image. Using the natural image database of over 20,000 images annotated with object segmentation masks (data curated by Microsoft and further annotated and analyzed in Rosenthal et al. 2018), we computed the object probability for each color using the procedure described in Rosenthal et al.^12^ as follows: (i) Each image is first encoded in the cylindrical representation of the Lu’v’ chromaticity space, namely the Hue-Chroma-Luminance color space (ii) Number of natural object and background pixels that fall within each color bin (from 240 colors bins at 24 equally spaced hue and 10 equally space chromas values) are then computed separately using the segmentation masks of natural objects. (ii) The object probability of each color is then derived as the number of pixels having that color in natural objects divided by the number of pixels having the same color in either natural objects or background. Once the probabilities are estimated, we compute the mean object color probability for each NSD image as the average of the probability over all color bins weighted by the number of pixels in the image that fall within each color bin.

**Texture**

We use entropy as a loose local statistical measure for texture. Entropy (E) is computed as the Shannon’s entropy of the grayscaled version of every image.

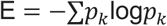

where *p_k_* is the probability of pixels to have a grayscale intensity value of k.

**Curvature index**

We used an image-computable curvature index to estimate the average curvature of contours in every image (as implemented by Li and Bonner (2020)^56^). This model convolves the grayscale version of each stimulus with a curvature filter bank with 176 different filters (16 orientations and 11 levels). Each filter in this bank functions as a curved contour detector with a specific orientation and curvature level. The grayscale image is also fed to an edge detection algorithm to find the edge pixels in each image. The overall curvature index is finally estimated by taking the average curvature over all the edge pixels in the image.

#### Quantifying the relationship between image-computable properties and Component 3 responses

##### Correlation analysis

We measured the relationship between each of the above variables and the responses of Component 3 to the shared image set using Pearson’s correlation coefficient.

##### Partial correlation analysis

We also performed a partial correlation analysis to assess the unique variance explained by each of the above image-level metrics in the responses of Component 3. We computed the correlation of the residuals resulting from a linear regression of all the above variables individually and food-salience ratings on the responses of Component 3 (the food-selective component). For food, we partialled out the effect of all the above confounders while computing the partial correlation.

II. Analysis on computationally matched food/non-food image pairs:

We identified pairs of images (one food and one nonfood) from the 5,445-10,000 image set viewed by each participant that produce similar activations in the last convolutional layer (‘*conv5*’) of an AlexNet^13^ pre-trained on ImageNet^16^ categorization. This matching analysis was performed on the entire image set of each participant at the subject-level (not just shared images), since having a larger stimulus set increases the chance of finding a stronger control pair. This computational matching procedure yielded 20-40 image pairs per participant. Mean food-nonfood similarity score, computed as the correlation between the features in the conv5 layer for the food image and the corresponding non-food image, across all pairs was thus very high (mean = 0.55, s.d. = 0.09); importantly, all pairs had a similarity score > 0.33. This was further substantially greater than the pairwise similarity among all the food images (mean = 0.16, s.d. = 0.17) and non-food images separately (mean = 0.11, s.d. = 0.18). Visual inspection further confirmed that these matched food-nonfood images contain similar visual features like similar colors and textures (example pairs shown in Figure 3C). We then performed a paired t-test to compare the responses of Component 3 to these food-nonfood pairs, separately for each participant.

III. Analysis on warm-colored non-food and cool-colored food stimuli:

Since the partial correlation analysis revealed a low, yet significant correlation of Component 3 responses with the object color probability measure (which reflects the warm-cool color continuum), we performed a subsequent analysis by directly pitting food preference against warm-color preferences. We sampled 50 food images from the lowest end of the object-color probability distribution over all 5,445-10,000 images (bottom 15 percentile) per participant and sampled non-food stimuli from the highest end of this distribution (top 15 percentile). This resulted in cool-colored food stimuli and warm-colored non-food stimuli. We then compared the responses of Component 3 to these two sampled subsets using an unpaired t-test separately for each participant. Example food and non-food stimuli from this selectively sampled distribution for one subject are shown in Figure 3D along with the distribution of the selection measure (object-color probability) for food and non-food stimuli.

While the food images from this analysis visually appeared to be cool-colored, this sampling procedure, however, could result in images where the ‘food’ itself is warm-colored since we are computing the mean object-color probability across the entire image. We thus conducted a subsequent analysis where we sampled 50 food stimuli such that the mean object-color probability over just the food pixels (as defined using the food segmentation masks obtained from MS-COCO annotations^52^) was at the lower end of this selection measure. This yielded images where the food itself was cool-colored. We repeated the statistical analysis by comparing the responses of Component 3 to these two sampled subsets and again found that the Component 3 responds much more strongly to food than non-food stimuli (Figure S5,B). This strongly suggests that the food-selectivity of Component 3 overrides any selectivity for warmer colors.

IV. Analysis on diverse subsets of food and non-food stimuli:

We selected diverse subsets of food and non-food stimuli that maximally span the representational space of different layers of a pre-trained DNN, such that the images within each subset are substantially more dissimilar (in terms of the average pairwise distance computed in the representational space of the corresponding layer) than what would be expected if the images were drawn at random from the respective set. The images were selected greedily so as to maximize the distance of each image with its closest neighbor. The procedure is outlined as follows: for each layer l,

i. we first randomly sample a food image,
ii. we then select the next image from the set of all food images viewed by each participant (N=10,000 in Phase 1 and 5,445 in Phase 2) as the image which has the largest correlation distance (1-r) to its closest neighbor among the already selected food images, where the distance is computed between the image features extracted at layer *l* and
iii. we repeat (ii) until we get the desired number of images (N=50).

The same procedure is repeated for the set of non-food images as well to get N=50 non-food images. These selected images are so diverse that a linear classifier trained to discriminate between these food and nonfood images using the features of the corresponding layer l performs at chance (never exceeding 53% across all layers), presumably because there is no remaining simpler visual characteristic shared by stimuli within the two subsets that a classifier can latch onto. We then compared the responses of Component 3 to these two stimulus subsets using an unpaired t-test, separately for each layer and each participant. This helps us address whether food selectivity is driven by only certain kinds of food images, which would indicate that it is not ‘food’ selectivity per se but rather a more restricted notion that applies to only specific instances of food; or whether the selectivity even persists under conditions of wide visual variability within food and within non-food images, which would in turn indicate that it is indeed ‘food’ selectivity construed more broadly.

V. Measuring the relationship between affective features and responses of Component 3:

Valence and arousal ratings were obtained from the NSD Meadows behavioral dataset which was released along with the NSD dataset (Further details provided in Allen et al., 2021^1^). These ratings were released for a total of 100 images from the shared image set viewed by all NSD participants. On this image set, we computed the correlation between the responses of Component 3 (averaged across subjects) and subject-averaged valence and subject-averaged arousal ratings separately. For fair comparison, we also report the correlation between food salience ratings and Component 3 responses on this small subset. The statistical significance of these correlations is assessed by computing the p-value of the obtained sample correlation coefficient for the null hypothesis of uncorrelation under the assumptions of a bivariate normal distribution.

### Analysis on curated Natural Scenes Dataset

To test whether a high proportion of exemplars of any category in the dataset might be sufficient for a component to emerge that responds selectively to that category, we sampled a subset of images with a fixed number of examples from each of 9 categories. These categories were selected because there were enough images in each subject belonging to those categories and include the following: face, food, clock, airplane, elephant, giraffe, horse, truck, motorcycle. Importantly, we chose an equal number of stimuli for each of these categories within each subject in this subset (although this number varied slightly across subjects because each subject saw different images, N=197, 207, 182, 191 per category for Phase 1 participants 1, 2, 5 and 7, respectively). The food images in this subset were drawn so that they are maximally heterogenous (dissimilar amongst themselves) following the procedure for sampling diverse subsets described above. The question was whether we’d get equally selective components for other categories that are in the same proportion as faces and food, which might suggest that the food-selective component could arise as an artifact of the data bias in NSD. These categories are also more visually homogeneous than food (e.g. airplane). We quantified the within-category visual similarity of images by computing mean pairwise correlations between the corresponding image features in the last convolutional layer of a pre-trained CNN (layer conv5 of AlexNet trained on image categorization using ImageNet). Distances computed in the feature space of trained DNNs (versus image space) are known to correspond well to perceptual image similarity measures, and are widely used as “perceptual distance” metrics^57, 58^; this metric is further also well-suited to capture similarities in mid-level visual features like texture; thus, the average pairwise image distance metric computed in this deep visual representational space for each category is likely to capture the perceptual homogeneity of that category (at least, as represented in the NSD).

We repeated the Bayesian NMF analysis on this curated dataset. On this subset, the BIC criterion suggested 7 instead of 20 components in each participant. We computed the selectivity of resulting components for each of the 9 categories using two indices: (i) Correlation (Pearson’s R), where we computed the correlation of component responses to the curated stimuli with a binary vector indicating whether the category was present/absent in the image over all stimuli and (ii) t-statistic, comparing the mean responses of the component to the category in question, versus all other stimuli. For each category, we then computed the maximum selectivity value based on either of the above indices over all 7 components, as reported in Figure 5.

The top 2 components (based on their highest correlation with any of these category labels) in each participant were still faces and food respectively. And the highest correlation of each remaining category with all the components was substantially lower than the selectivity of the top 2 components for faces and food, respectively. This control analysis indicated that data bias (either a large number of food images in NSD or some form of visual homogeneity among the food images within NSD) cannot explain the existence of the food-selective component.

### Analysis on the independent BOLD5000 dataset

We assessed whether food-selective responses can also be identified in other independent datasets beyond NSD. To test this, we analyzed another publicly available large-scale dataset, namely, the BOLD5000 dataset. This dataset comprised BOLD responses from four participants (CSI1, CSI2, CSI3, CSI4), while they each viewed several thousands of natural images, though most images had only single repetitions. Importantly, the shared set of 1,000 images viewed by all Phase I NSD participants were also viewed by all BOLD5000 subjects, with the exception of subject CSI4 who only viewed 594 shared images. We used these overlapping images to localize the food component in the ventral visual stream of subjects CSI1-4. This localization procedure relies on inferring the voxel weights corresponding to Component 3 (the food-selective component) in new participants using the response profile of Component 3 derived from NSD over the overlapping image set. In the non-negative matrix factorization parlance, this amounts to inferring only one weight matrix (the components by voxel weight matrix), when the other matrix (the response profile matrix) is known, subject to non-negativity constraints. Here, the latter is fixed to the Component 3 responses averaged across NSD subjects for the overlapping image set (N=1,000 for CSI1-3 and N=594 for CSI4). Mathematically, given component responses to N overlapping images derived from NSD as **R** *(Nx1),* and the data matrix **D** (N x V) containing the responses of all V voxels to these N stimuli in a BOLD5000 subject, the non-negative voxel weights **W** (Vx1) for the component can be estimated by minimizing the expression,

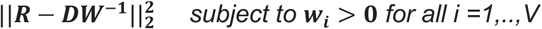

This optimization problem is convex and the optimal voxel weights **Ŵ** can be derived following the standard routine for solving the non-negative least squares problem based on the active set algorithm^59^. With these inferred voxel weights, we can then estimate the component responses to novel stimuli unique to the BOLD5000 dataset (**D_U_**) as follows,

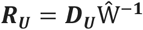

We restrict our focus to stimuli that had food/no-food annotations in the BOLD5000 dataset, namely, the MS-COCO images. We excluded all MS-COCO images that were viewed by any of the NSD participants from this analysis. We then computed the food-selectivity of estimated component responses to these stimuli as the correlation (Pearson’s R) between component responses and a binary vector indicating whether the image contained food or not. These images were further rank-ordered by their response magnitude and colored by food labels for ease of response visualization. We further performed a control analysis by running the component localizer in other areas of the visual cortex, including early visual areas as all intermediate and high-level lateral and parietal areas, and computing the food-selectivity of the estimated component in each case. These ROIs, including the ventral visual stream ROI as used above, were defined by co-registering the *streams atlas* from NSD to the BOLD5000 anatomical space.

### Quantification and statistical analysis on all component voxel weights

#### I.#Agreement with functional localizer statistics

Once the voxel weights are projected back into anatomical coordinates (in the native space of each NSD participant), we can also compute the quantitative agreement between these voxel weights and the voxel-level selectivity for different categories as estimated with the independent functional localizer runs in NSD (fLOC). We computed the correlation between the voxel weights of each component against the voxel-wise t-statistic of the component’s preferred category as obtained with the fLOC experiments by contrasting responses to each category against all other stimuli. For e.g. the face component voxel weights were correlated against the t-value contrasts for responses to the domain of faces over responses to all other stimuli. Note that food was not defined as a domain in the NSD fLOC experiment, since a selectivity for food in the visual cortex had never been described before; thus, we cannot perform a similar analysis for Component 3.

#### II.#Anatomical similarity between saturation-responsive visual cortex and Component 3

The anatomical distribution of Component 3 appeared to overlap with previously studied color-biased regions^60^. To quantify the similarity between the anatomy of saturation-responsive regions and Component 3, we conducted a subsequent analysis. We first extracted non-food images (food salience rating of zero) from the shared set of 515 images viewed by all participants. We next computed the correlation between the saturation of all non-food stimuli (N=356) and the responses of all VVC voxels to the corresponding stimuli in order to construct a saturation-responsive voxel weight map per participant. Relationship between food selectivity and saturation-responsiveness in the ventral visual pathway is finally assessed by correlating this saturation-responsive weight map with the voxel weight map for each of the 5 components. These correlations were transformed to z-scores using Fisher’s z-transformation for statistical comparisons.

#### III.#Quantifying the spread of voxel weights per component

We further characterized the distribution of voxel weights for each component using quantitative measures of sparseness and statistical measures of skewness and kurtosis.

(a) **Skewness** of the voxel weight distribution ***w***_***c***_ for each component *c* is computed using the Fisher-Pearson coefficient of skewness, calculated as:

Skewness 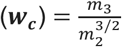,

where *m*_2_ and *m*_3_ are respectively the second and third sample central moments of the voxel weights ***w***_***c***_ for each component *c*. The *r*th sample moment *m_r_* are computed using the standard formula as, 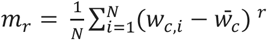

where *w_c,i_* is the voxel weight for component c in voxel i and 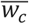 is the mean component weight across all N voxels. This measure is computed separately for the voxel weights per component and per participant where different participants have differing numbers of voxels (N∼6,500-9,000).

For a gaussian distribution (perfect symmetry), the skewness is zero; positive values indicate a rightward skew with more voxels that have higher weights on the component whereas negative values point towards a leftward skew.

(b) **Sparseness** in the voxel weights of each component, ***w***_***c***_ with N voxels is computed using the definition of [Hoyer et al., 2004] as,

Sparseness 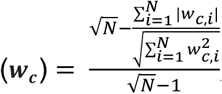,

Sparseness is 1 when only a single voxel has a non-zero weight on the component and is zero when all voxel weights are equal (non-sparse distribution). Values between 0 and 1 indicate intermediate levels of sparsity, interpolating smoothly between the two extremes.

(c) (Excess) **Kurtosis** of the voxel weight distribution for each component *c* is computed following Fisher’s definition, as the ratio of the fourth sample central moment of the voxel weights ***w***_***c***_ and their second central moment squared,

Kurtosis 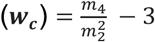,

Here, 3 is subtracted to provide a simple comparison to the Gaussian distribution which yields a kurtosis of zero under the above definition. Higher values (above 0) indicate a super-gaussian or heavy-tailed distribution indicative of sparsity.

#### IV.#Quantifying the inter-subject heterogeneity of voxel weights per component

We transformed the voxel weight maps from the native space of each participant to a common anatomical space (MNI 1mm) in order to measure inter-subject alignment in the anatomy of each component. For each component, this alignment was measured using correlation (Pearson’s R) between the co-registered weight map of each participant and the average voxel weight map for that component across the other 7 participants (averaged across all 8 folds).

### Encoding model of the inferred components

We used a CLIP-ResNet50^15^ convolutional neural network (CNN) model to predict the response of the inferred components from the NMF analysis. The encoding model was designed to map the features from a given layer of the CNN model to the inferred responses from the component analyses (see Ratan Murty et al. (2021)^14^ for more details). Importantly, we fixed all the hyper-parameters of the model based on the data from Phase 1 subjects. Specifically, we fixed the model layer (block4-1-conv2). The model features corresponding to the images used in the experiment were extracted for this layer. Next we mapped the extracted features to the inferred component responses of Phase 2 subjects via a ten-fold regularized ridge-regression (the ridge parameter fixed at 0.01). Even though the model was trained on data from Phase 2 subjects, it was evaluated on data from Phase 1 subjects (thus cross-validating on both subjects and images). The model prediction accuracy was calculated as the Pearson correlation between the predicted response of the model (over folds) and the observed response. Our CLIP-ResNet50 encoding model is image-computable and can be used to predict the observed responses for images not included in the NSD. We obtained predictions for: 1) The large publicly available ImageNet dataset which has diverse stimuli from 1000 stimulus categories (N = 1,281,167 images). 2) Black and white versions of the same 1.2M images as in 1, 3) color and grayscale versions of the texture-matched Downing pairs. These images were previously used to test and reject the food selectivity hypothesis in the brain^17^. Predictions were obtained for both color and grayscale versions of these images. (Figure 4). 4) Handpicked images that were matched across a number of stimulus features. See Figure 4 for examples. Predictions were obtained for both color and grayscale versions of these images (Figure 4).

## Data availability

The NSD dataset is freely available at http://naturalscenesdataset.org. Images used for NSD were taken from the Common Objects in Context database (https://cocodataset.org).

## Supplementary Movies

Supplementary Movie 1: Top 25 images producing the highest response in each of the top 5 components in each participant (both Phase 1 and Phase 2). https://drive.google.com/file/d/15m9Ys_ougo0WN09KU1SyDnU8qAvfu_yV/view?usp=sharing

Supplementary Movie 2: Top 5% voxels with the highest weights on each component are visualized on cortical flatmaps for each of the 8 subjects. Established regions of interest, defined from the functional localizer scans by computing the contrast of preferred versus all other stimuli, are shown in white outlines (t-value > 2.5). Voxel weight maps of face, food and word components are also visualized together on the RGB colormap to show component overlap for each subject. https://drive.google.com/file/d/1jab7RFrUL0eA0CGDrU5VCjwyUYpjR6qs/view?usp=sharing

## Supplementary Figures

**Figure S1:**
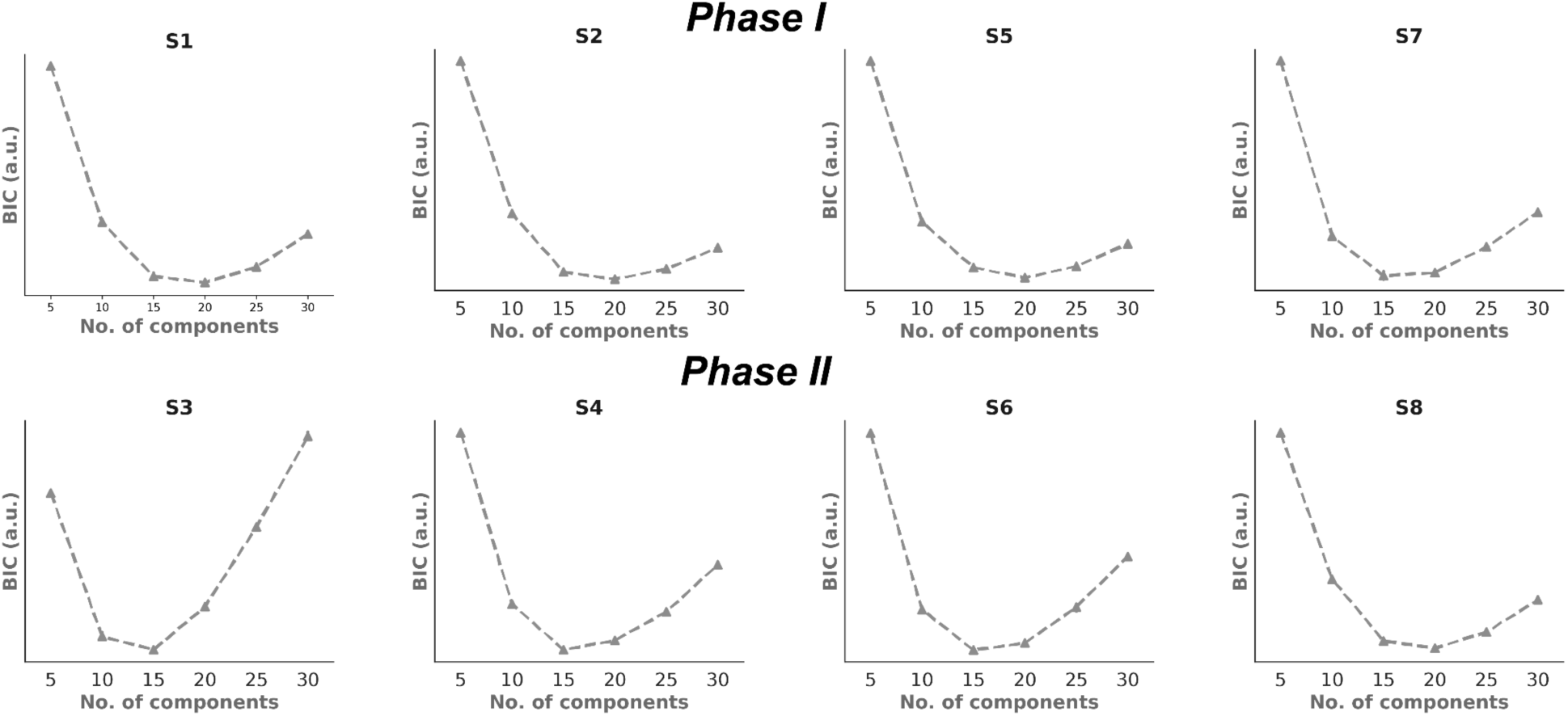
Bayesian Information Criterion (BIC) used for selecting the number of components. The optimal number of components, as suggested by the BIC, were 15-20 in all subjects.

**Figure S2:**
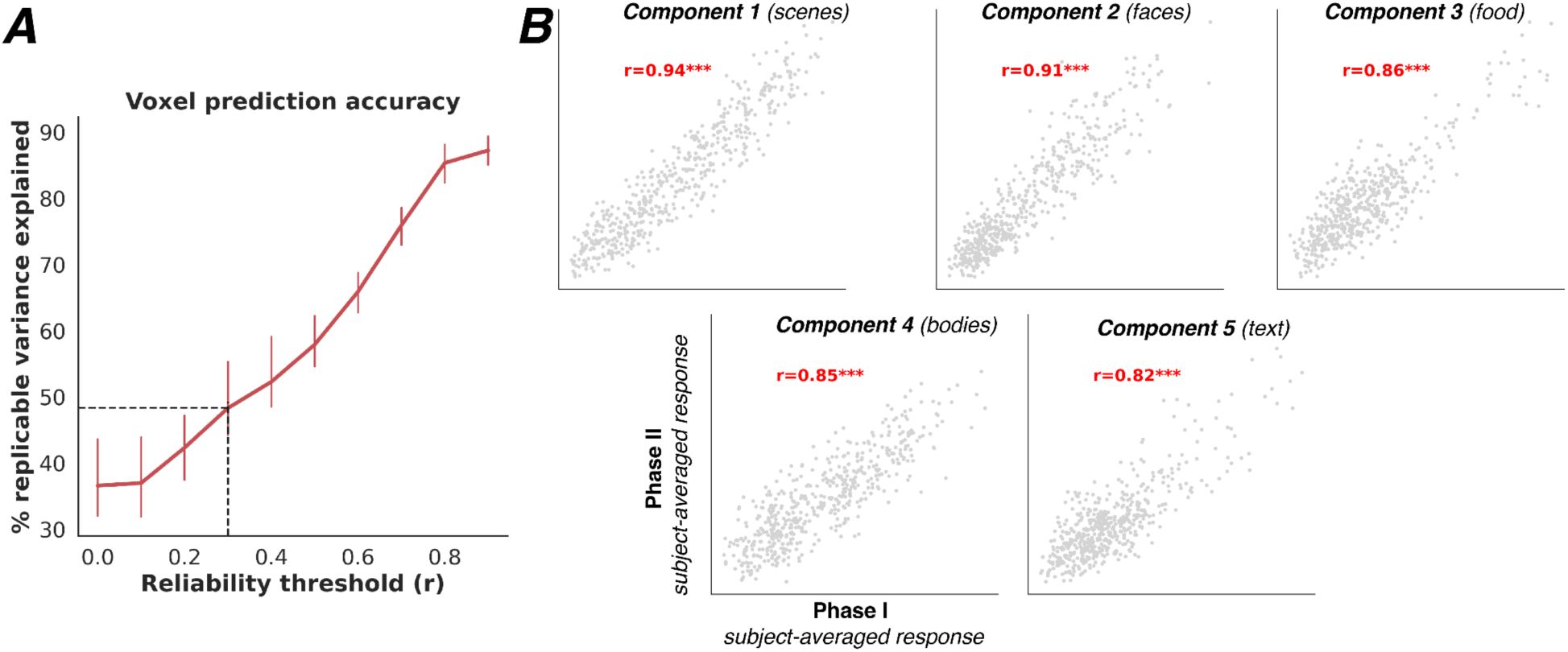
**A.** Accuracy of the 5 component model derived from Phase 1 participants in explaining responses of raw voxels in the ventral visual stream of held-out subjects from Phase 2. Accuracy is expressed as the % of replicable variance explained by the five components taken together (median across voxels) as we vary the reliability threshold for voxel selection (median noise ceiling expressed in correlation units) in accuracy computation. The five-component model can explain ∼50% of the replicable variance in the response of voxels with 0.3 or greater reliability (which includes about 46% voxels in the ventral visual stream averaged across different participants). **B**. Scatter plots showing the subject-averaged responses of each component in Phase I against the subject-averaged responses of the same component identified in Phase II.

**Figure S3:**
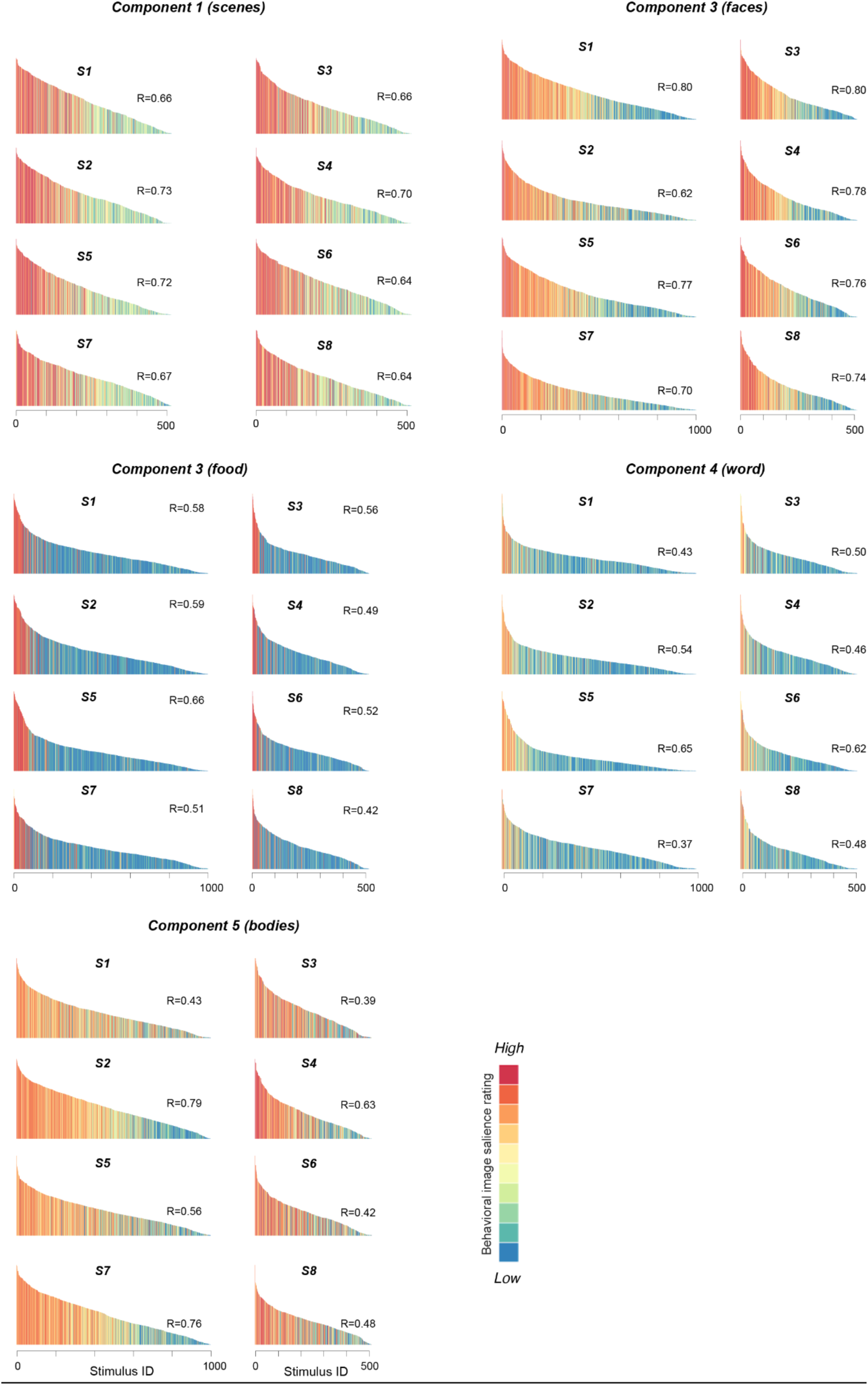
Response profile of each component in each individual across all shared images, sorted by the response magnitudes and colored by the salience rating of the preferred category for that component for every image. Left panel shows the response profiles of Phase 1 participants who each viewed a common set of 1,000 images whereas the right panel shows the results for Phase 2 participants who viewed 515 shared images, distinct from the images unique to each participant. For Component 1, responses are shown only for 515 images in Phase 1 participants since we obtained expert scene salience annotations for 515 images viewed by all 8 participants. Correlations between salience ratings for the preferred dimension for that component and subject-specific component responses across all shared images are displayed on the top right of each subplot.

**Figure S4.**
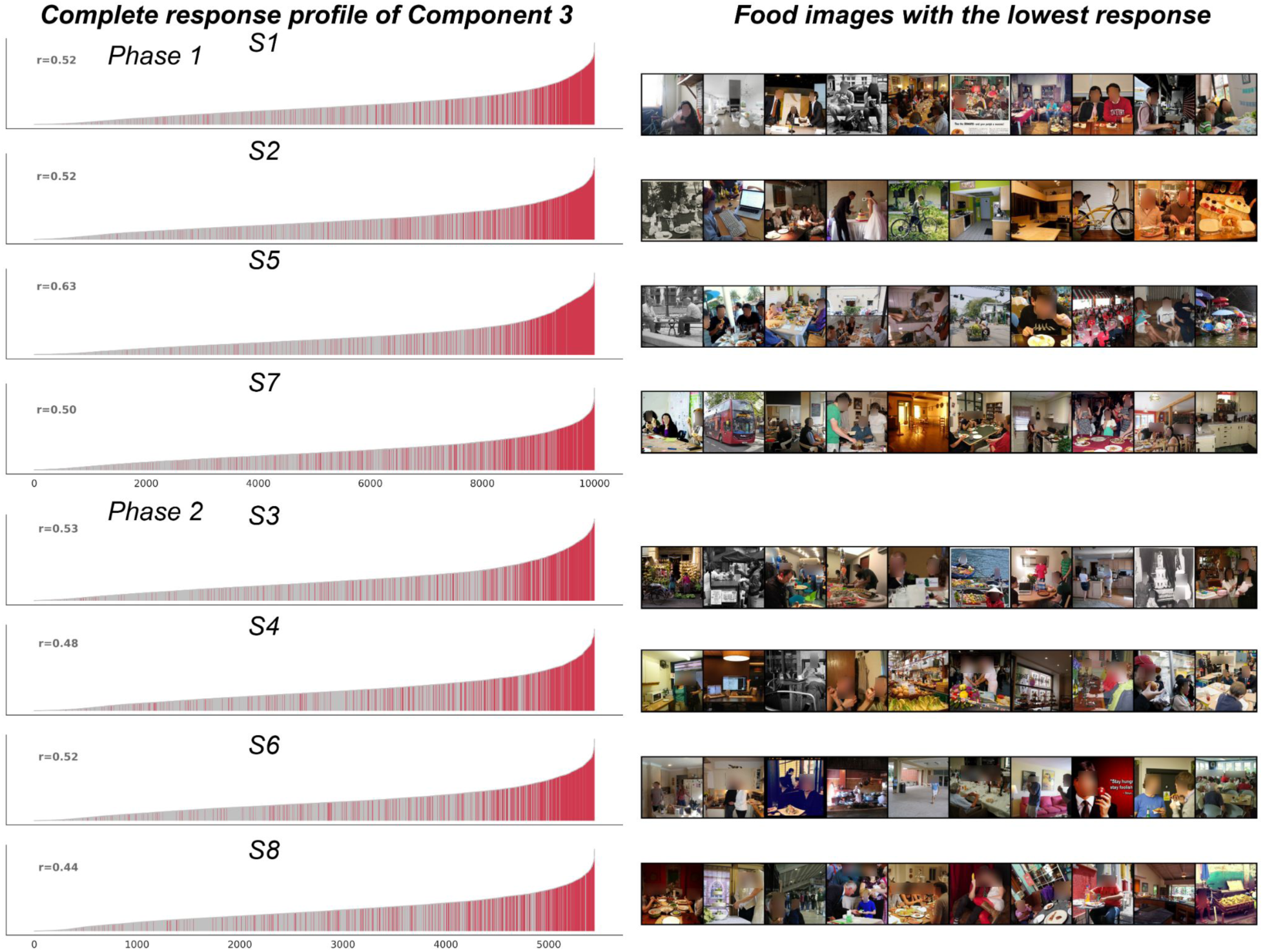
Complete response profile of the food component across all 10,000 (Phase One participants, A) or 5,445 (Phase Two participants, B) images, colored by food labels obtained from MS COCO annotations. Binary food labels (red) can explain a large proportion of response variation for this component in all participants (r=0.44-0.63). Right panel shows the images labeled as ‘food’ that produce the *lowest* response in the food component in each participant. As can be seen in the figure, most of these are images that were either mislabeled or where the food is not particularly salient.

**Figure S5:**
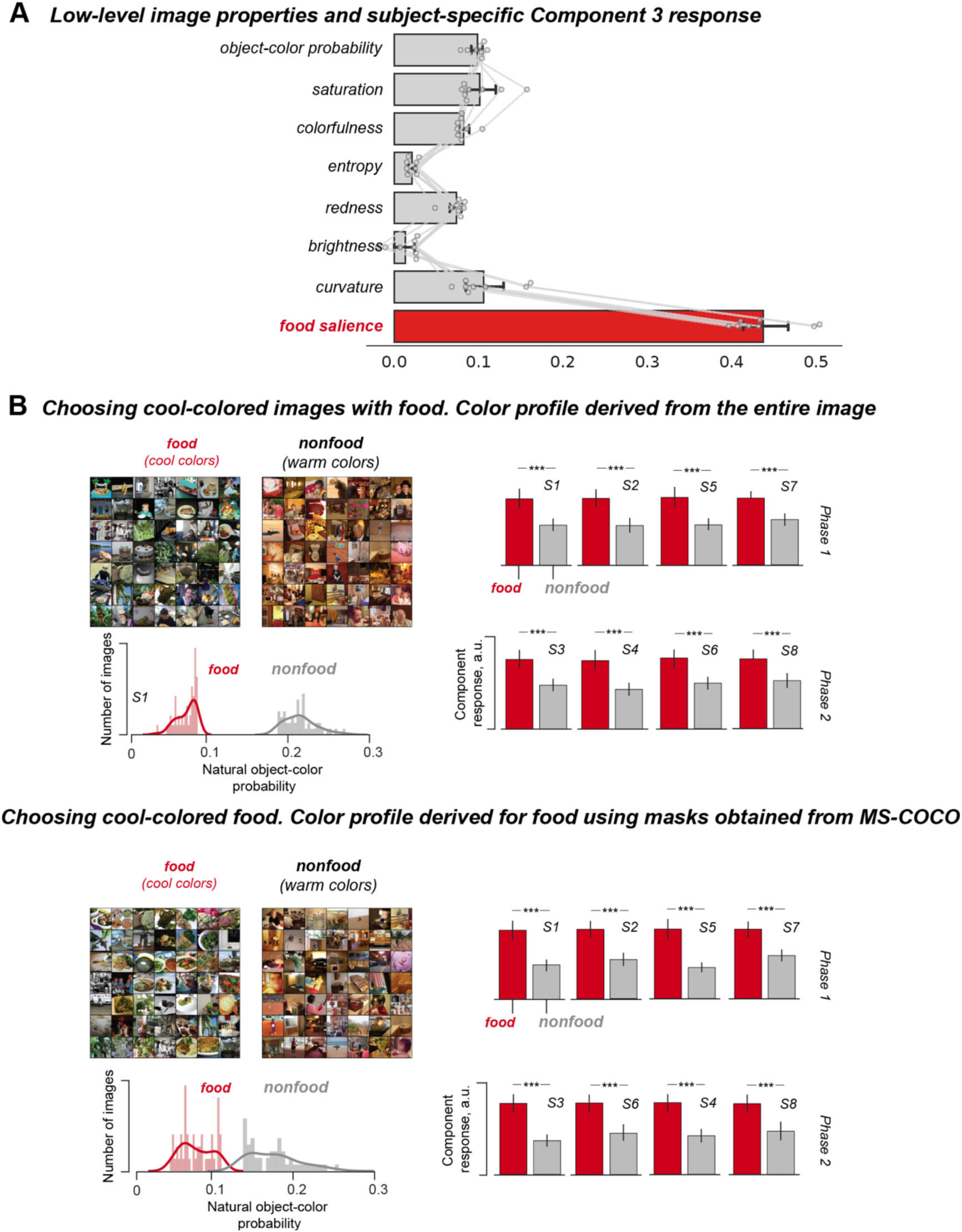
A.The partial correlation across stimuli between the magnitude of the Component 3 response in individual subjects and various image-computable feature dimensions (with the effect of food salience removed) and rated food salience (with the effect of all other feature dimensions removed). The partial correlation with food salience is significantly greater than any of the others (all ps < 0.00001). B. (Top) Response of the Component 3 in each participant to sets of stimuli chosen such that the images of food were very low and the nonfood images were very high on the object-color probability measure (Rosenthal et al., 2018). (Bottom) Same as the top except that sets of food stimuli were chosen such that the mean object-color probability across food pixels (segmentation masks for food were obtained using MS-COCO annotations) was very low.

**Figure S6:**
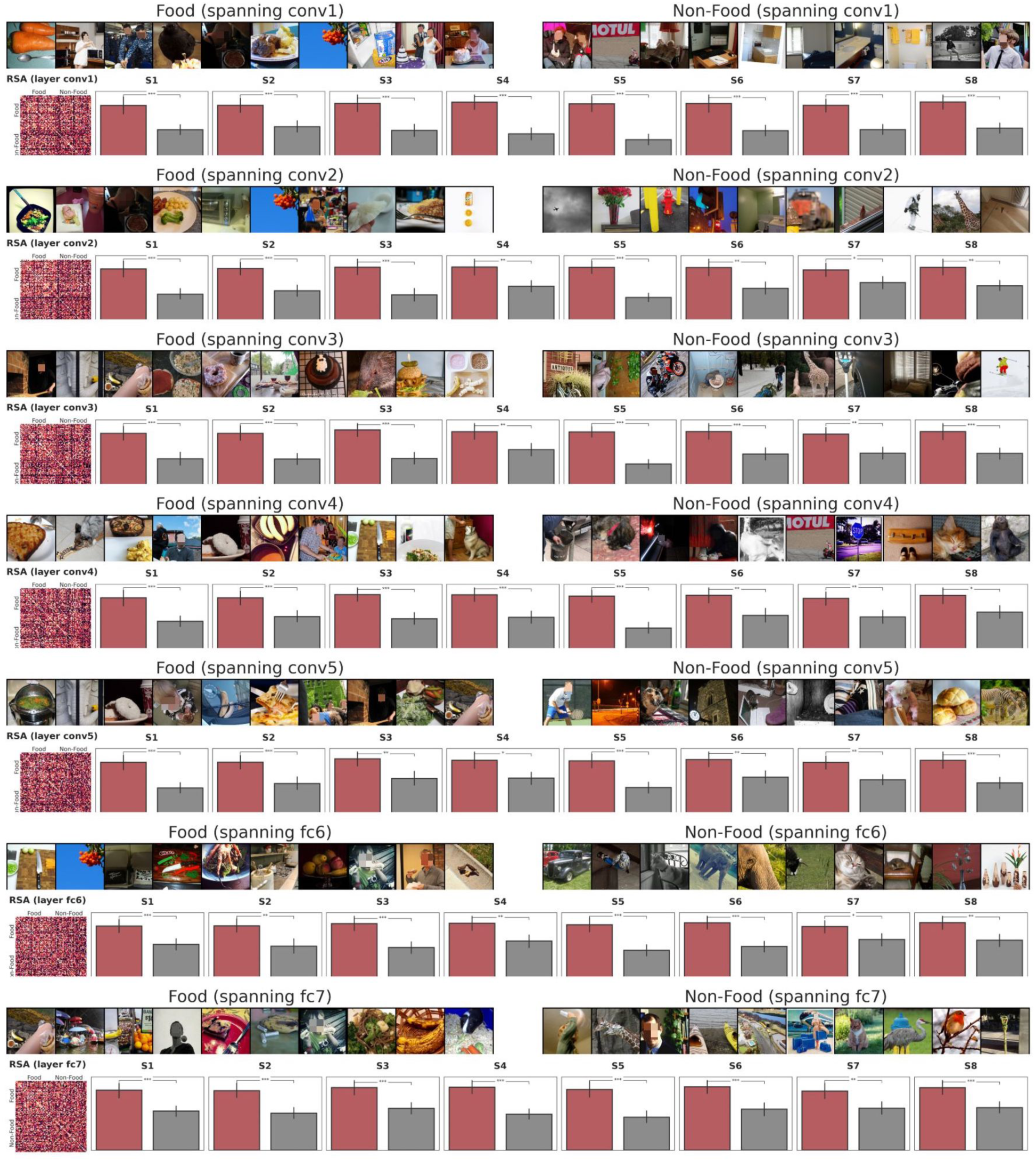
Comparing responses of the food component to diverse subsets of food and non-food stimuli. Subsets of food and non-food stimuli that maximally span the embedding space of different layers of an ImageNet-trained AlexNet are selected based on a greedy sampling procedure described in the Methods. These subsets are indiscriminable by the corresponding layer, as also suggested by the RSA plot for one subject (left). Examples of food and non-food stimuli from each subset are shown above the responses of the food component to the corresponding subsets. Error bars show the 95% confidence interval around the estimated mean. Response to the subset of food stimuli is significantly greater than the response to the subset of non-food stimuli in *all* cases.

**Figure S7.**
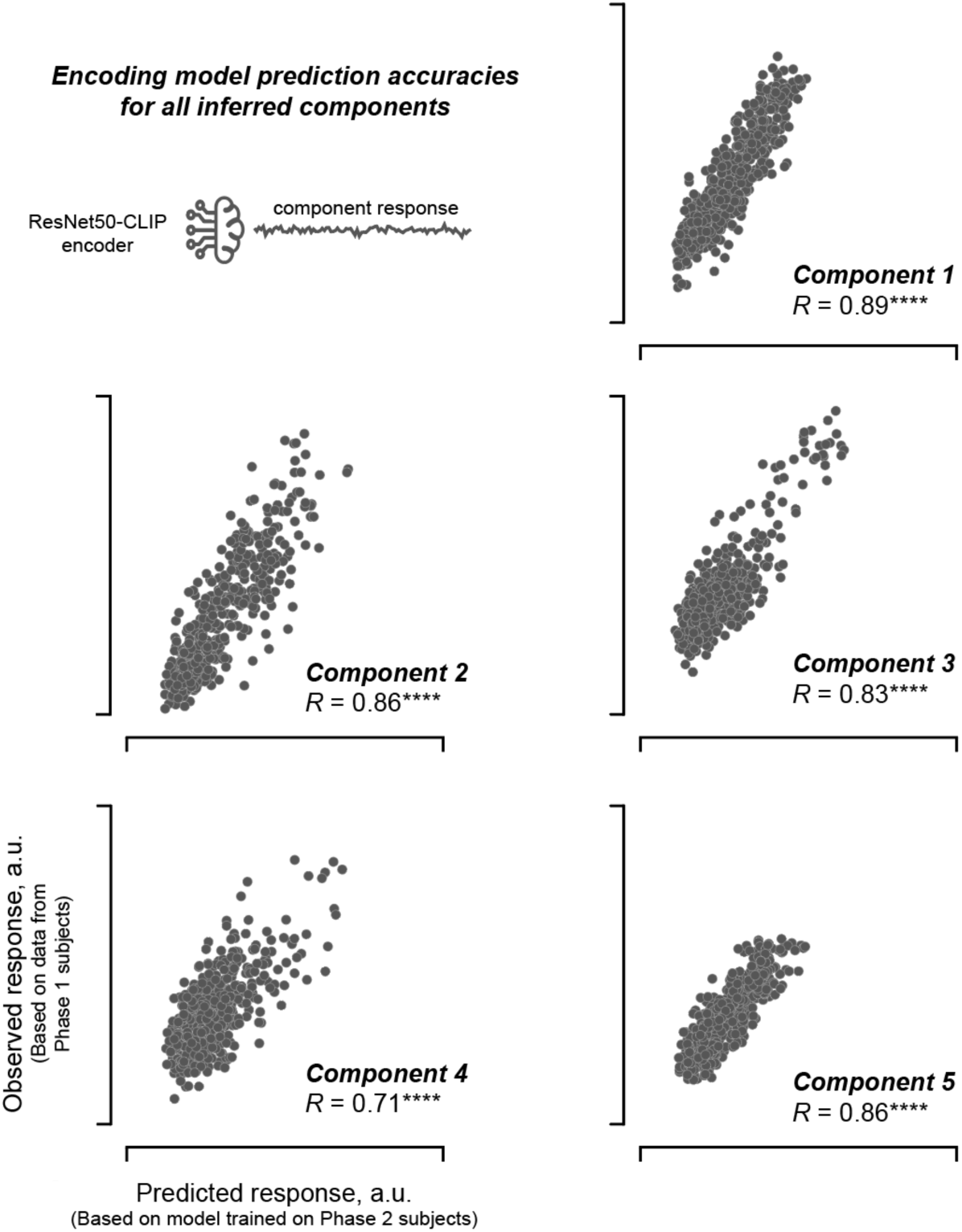
Predictive accuracy of a Resnet50-CLIP based encoding model. Scatterplot between the predicted response on the x-axis (based on model trained on data from Phase 2 subjects) and the observed response on the y-axis (based on observed data from Phase 1 subjects). Each dot is a given image and the predictions were always estimated on images not used in the model training procedure.

**Figure S8.**
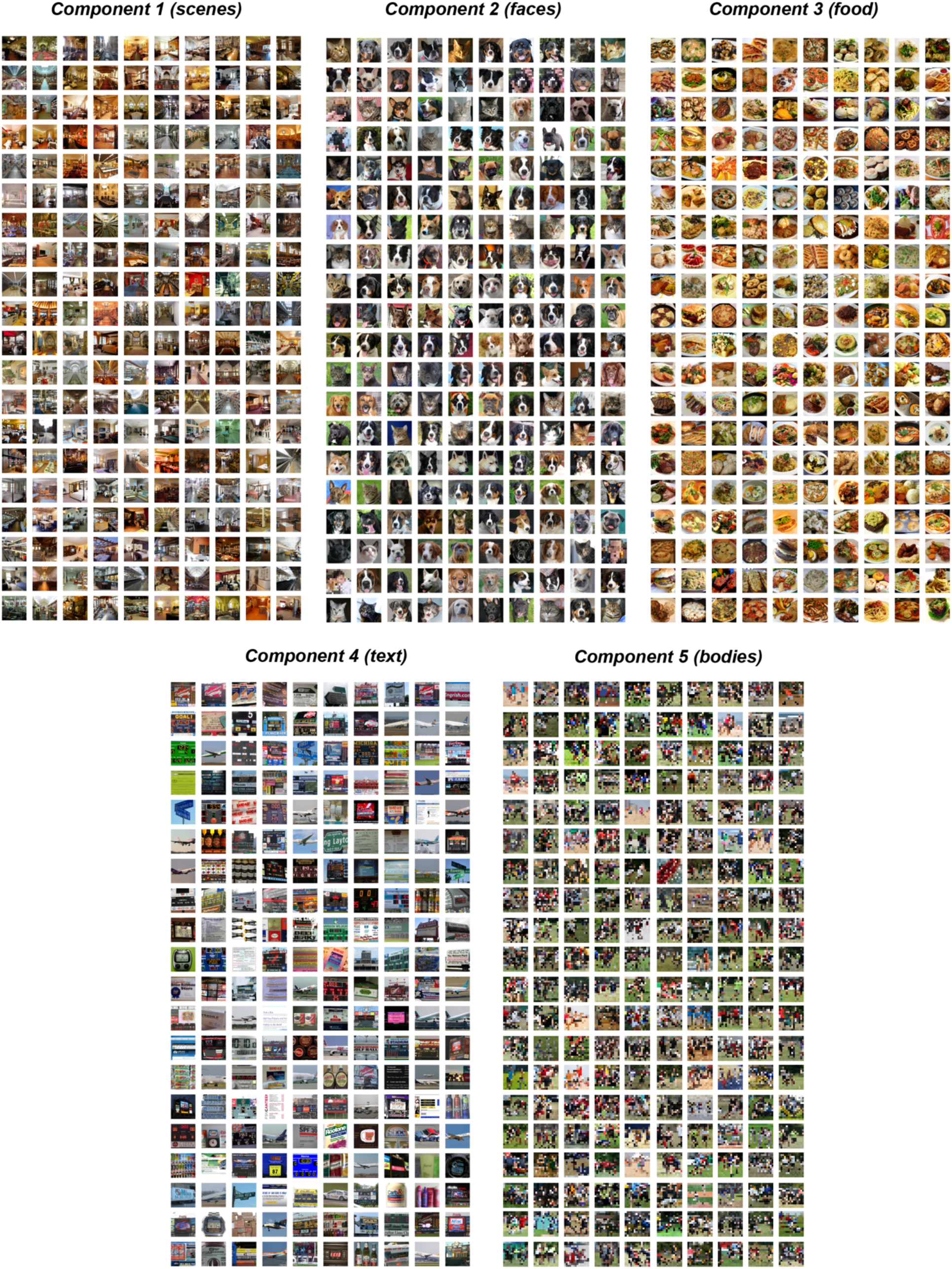
Top 200 images (among the ImageNet stimuli) predicted by the Resnet50-CLIP encoding model to activate each of the 5 components

**Figure S9:**
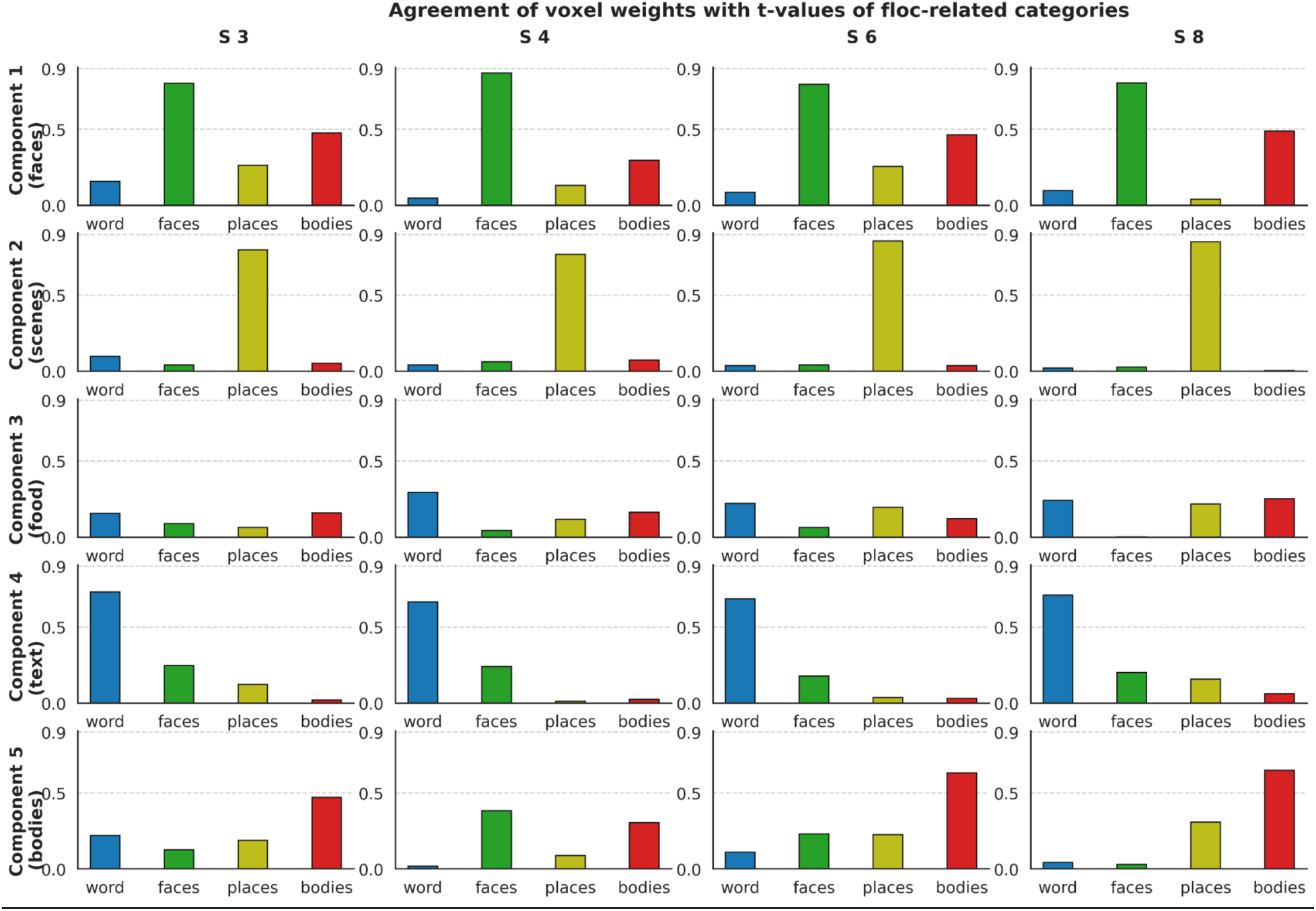
Weight maps for the faces, scenes, text and bodies weight maps agree with the corresponding contrast maps from the fMRI localizer scan. The correlation across voxels of the weight map for each component with the t value of the contrast from the fMRI localizer scan between words versus all other stimuli (blue), faces versus all other stimuli (green), places versus all other stimuli (yellow), and bodies versus all other stimuli (red). Stimulus categories in the floc experiment include words (characters), faces, places, bodies and objects.

**Figure S10:**
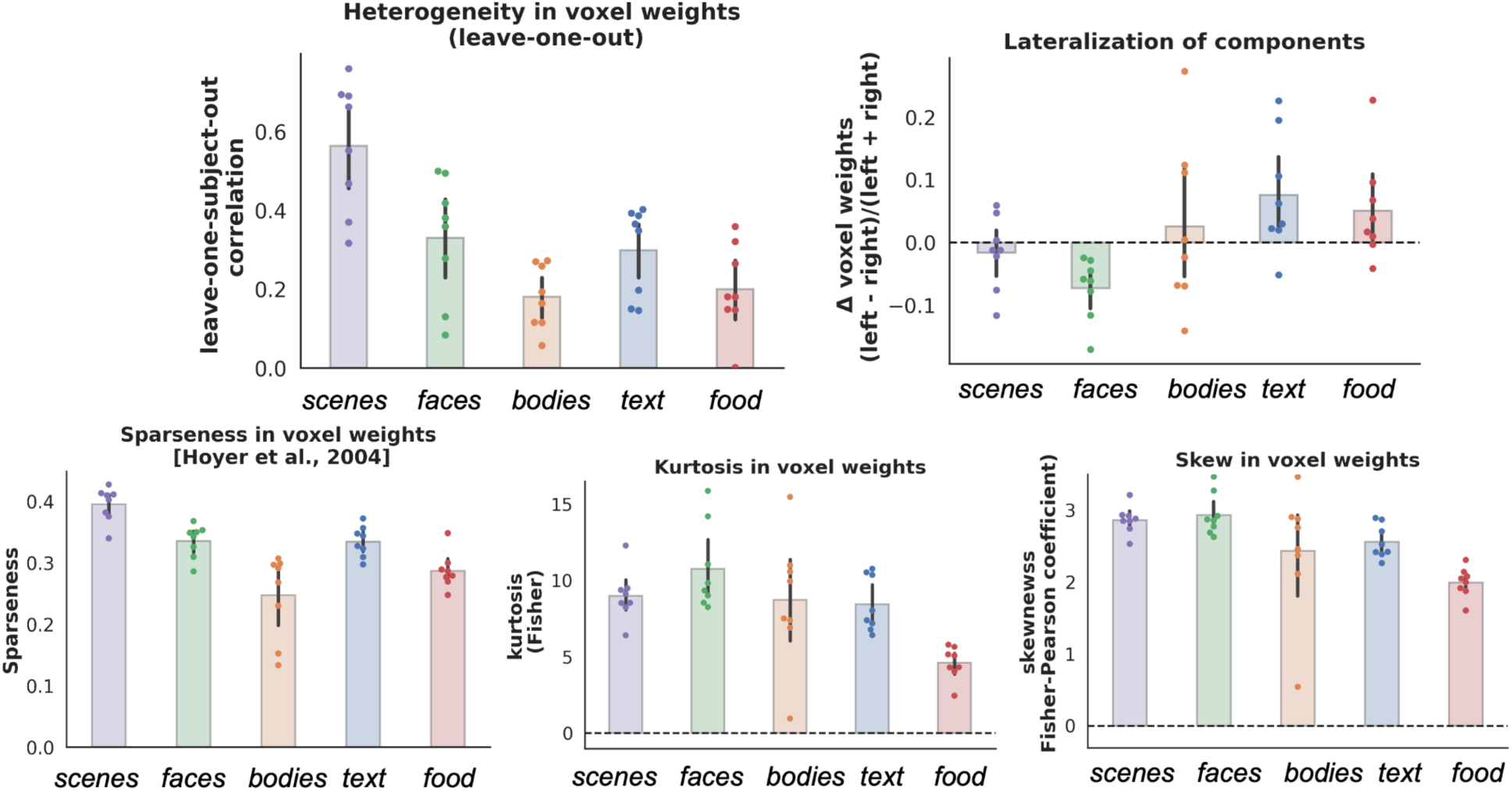
Quantitative<u> measures of voxel weight distributions for each component.</u> Upper Left: Correlation of the weight map for each component between each participant and the average weight map across the other 7 participants (averaged across all 8 folds) when co-registered into a common anatomical space (MNI). Upper right: Lateralization of components is computed as the difference between the average voxel weights in the left and right hemisphere divided by the sum of the two. A value of 1 indicates perfect left lateralization whereas negative values indicate right lateralization. Lower Left: A sparseness measure for the voxel weights of each component is computed based on the relationship between L1 and L2 norms following Hoyer et al. (2004); higher values above 0 indicate higher sparseness. Lower Middle and Lower Right depict the kurtosis and skewness of the voxel weight distributions of each component. The corresponding values for a Gaussian distribution (skew=0, excess kurtosis=0) are marked as dashed horizontal lines. All components have voxel weights that are positively skewed and kurtotic, relative to a Gaussian, indicating a peakier, heavy-tailed distribution skewed towards higher values. Each scatter point corresponds to a single subject.

**Figure S11:**
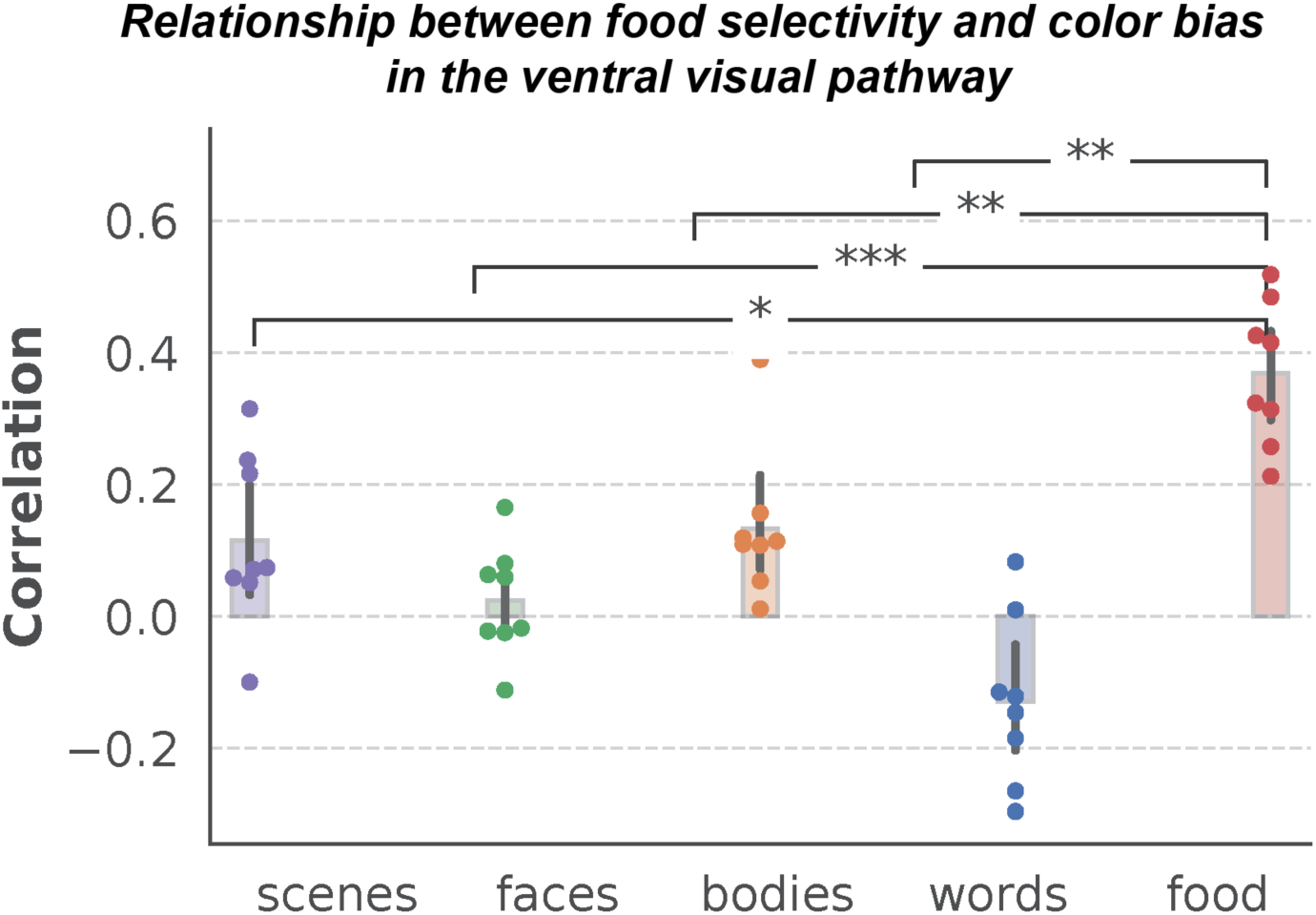
A color-bias (saturation-responsive) map in the VVC is computed for each subject by measuring the correlation of voxel responses to non-food stimuli with the saturation index of corresponding images. Relationship between food selectivity and color bias in the ventral visual pathway is assessed by correlating the color-bias map (computed on non-food stimuli) with the voxel weight map for each of 5 components. Each scatter point is an individual subject. These correlations were transformed to z-scores using Fisher’s z-transformation for statistical comparisons. The food component weight map exhibited a significantly higher correlation with the color bias map than any other component (p < 0.01 for all 4 comparisons using a paired t-test).

**Figure S12:**
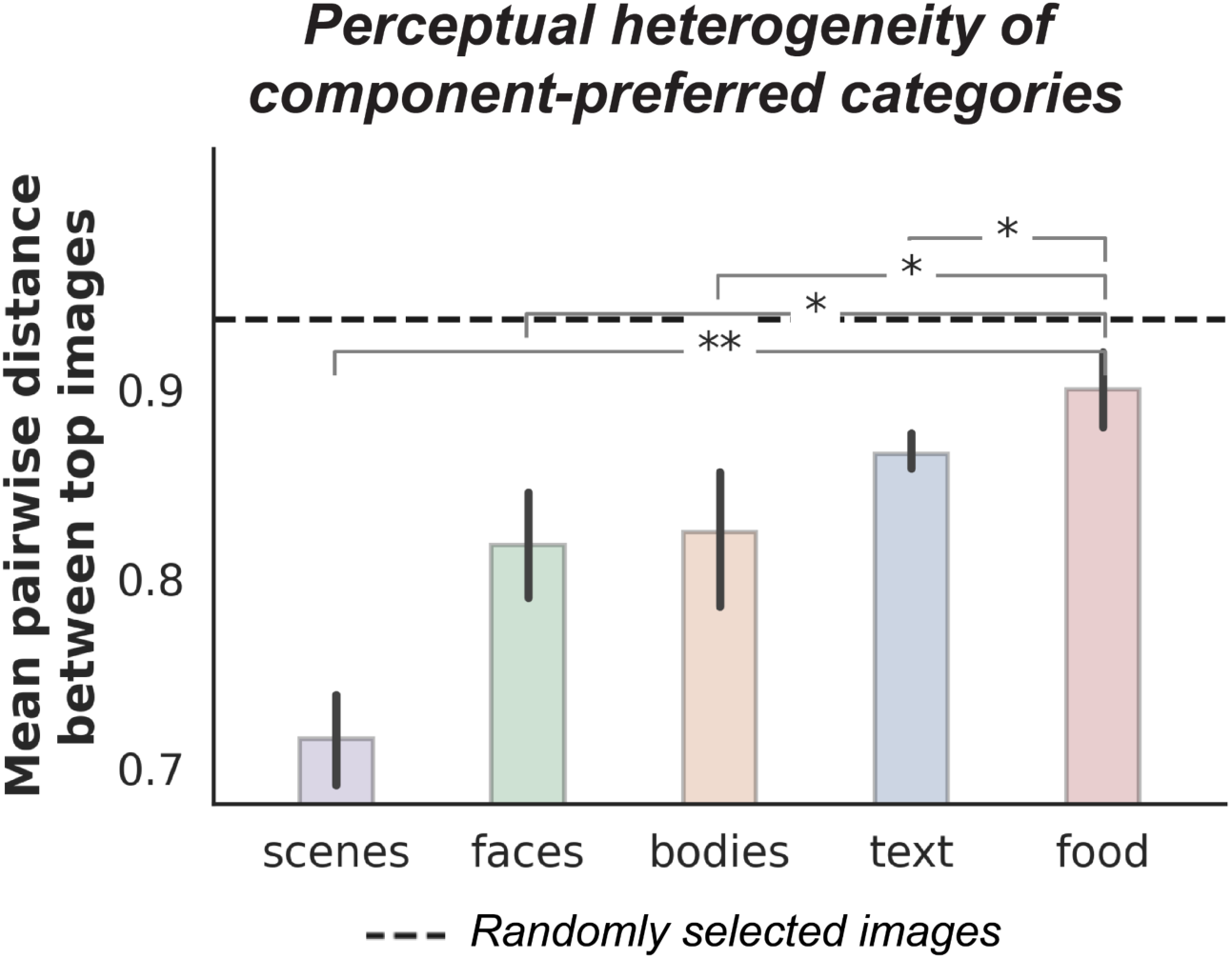
Perceptual heterogeneity of different categories in NSD is estimated by computing the average pairwise correlation distance (1-r) between the top 10% images (among the 5,445-10,000 images viewed by each participant) producing the highest response in each component in the embedding space of the last convolutional layer of AlexNet (‘*conv5*’). Dashed black line depicts the mean pairwise distance between the same number of randomly selected images. Error bars indicate the 95% confidence interval around the estimated mean distance across subjects.

